# 50-color phenotyping of the human immune system with in-depth assessment of T cells and dendritic cells

**DOI:** 10.1101/2023.12.14.571745

**Authors:** Andrew J. Konecny, Peter Mage, Aaron J. Tyznik, Martin Prlic, Florian Mair

## Abstract

We report the development of an optimized 50-color spectral flow cytometry panel designed for the in-depth analysis of the immune system in human blood and tissues, with the goal of maximizing the amount of information that can be collected using currently available flow cytometry platforms. We established and tested this panel using peripheral blood mononuclear cells (PBMCs), but included CD45 to enable its use for the analysis of human tissue samples. The panel contains lineage markers for all major immune cell subsets, and an extensive set of phenotyping markers focused on the activation and differentiation status of the T cell and dendritic cell (DC) compartment.

We outline the biological insight that can be gained from the simultaneous measurement of such a large number of proteins and propose that this approach provides a unique opportunity for the comprehensive exploration of the immune status in tissue biopsies and other human samples with a limited number of cells. Of note, we tested the panel to be compatible with cell sorting for further downstream applications. Furthermore, to facilitate the wide-spread implementation of such a panel across different cohorts and samples, we established a trimmed-down 45-color version which can be used with different spectral cytometry platforms.

Finally, to generate this panel, we utilized not only existing panel design guidelines, but also developed new metrics to systematically identify the optimal combination of 50 fluorochromes and evaluate fluorochrome-specific resolution in the context of a 50-color unmixing matrix.

## Background

The immune system serves as the body’s defense system against pathogens and is also essential for maintaining steady-state homeostasis in tissues (1) and preventing the development of malignant tumors (2). The composition and activation status of immune cells in the periphery and in tissues can be used to extrapolate immune cell differentiation and function. To facilitate data interpretation, an immune cell population is ideally analyzed in the context of other immune cell populations. Thus, to comprehensively study the functional state of the immune system, it is highly beneficial to capture as much information from as many different cell types as feasible. This is particularly relevant for assessing immune cell function *in situ*, for example in human tissue samples (3). Moreover, these human tissues are often limited in size and availability, e.g. tissue biopsies (4) or resected pieces of tumor tissue (5), which precludes parallel analysis with multiple panels or multiple applications. The development of an analysis approach that can provide broad and, for some cell subsets, in depth-phenotyping, paired with the ability to preserve cell populations of interest for downstream applications such as single-cell RNA-sequencing, is hence of importance.

The interactions between professional antigen-presenting cells (APCs) and different T cell subsets (6) are of particular interest in the context of studying anti-tumor immune responses (7). Dendritic cells (DCs) are highly specialized APCs, and generally divided into cross-presenting cDC1s and cDC2s (8), with CD163^+^ cDC3s being a more recently described subset during inflammatory conditions (9). Each of these DC populations appears to have a distinct function for steering an adaptive immune response. T cells consist of conventional CD4^+^ T cells, regulatory CD4^+^ T cells (Tregs), CD8^+^ T cells, γδ T cells and subsets of T cells with semi-invariant T cell receptors including mucosal-associated invariant T cells (MAIT cells) and invariant NK-T cells (10).

The panel presented here (Figure 1A) was designed to comprehensively capture the differentiation and activation status of T cells and APCs, while also measuring B cell, NK cell and innate lymphoid cells (ILCs) phenotypes (list of markers depicted in Figure 1B and Table 2). Optimization was done on human cryopreserved PBMCs, but the panel includes CD45 as a pan-hematopoietic marker and has been tested on human tissue-derived leukocytes (data not shown).

**Figure 1:**
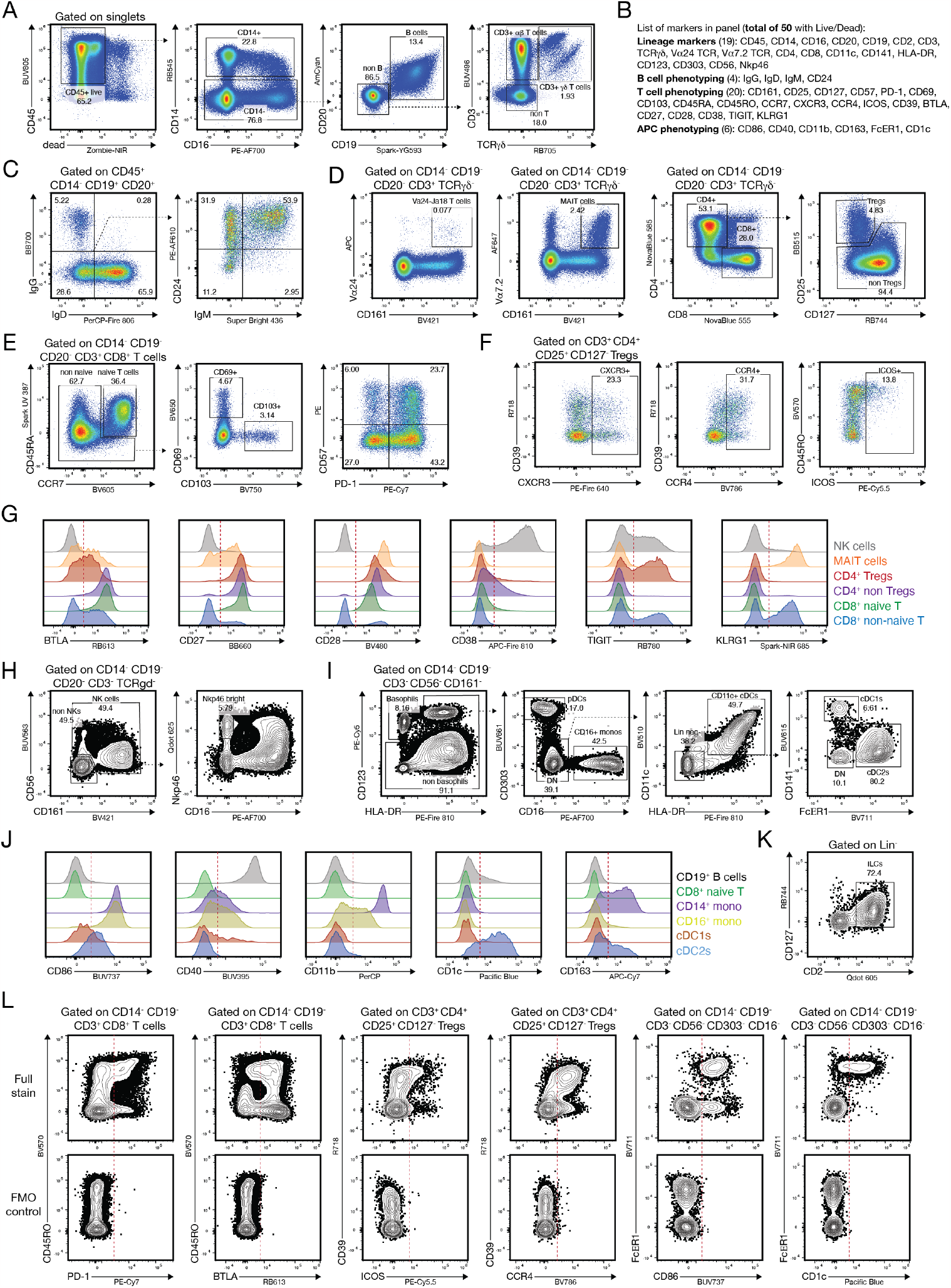
Overview gating of the 50-color panel on cryopreserved PBMCs. PBMCs were obtained from commercial vendors, stained as described in the online section of the manuscript and acquired on a BD FACSDiscover™ S8. The optical configuration of the instrument is described in Online Figures 1-2. Additional gating and staining controls are shown in Online Figures 9-11. Pre-gating of plots is annotated in the figure or indicated by dotted black arrows. For some plots different donors are shown for clarity. The gating strategy has been devised in such a way that the staining pattern for every marker in the panel can be shown at least once on a single A4 page. The raw data has been deposited on Flowrepository with the identifier FR-FCM-Z73V. Abbreviations: BUV: Brilliant Ultraviolet; BV: Brilliant Violet; BB: Brilliant Blue; RB: RealBlue; AF: AlexaFluor; PE: Phycoerythrin; APC: Allophycocyanin; Qdot: Quantum Dot; NIR: Near-Infrared; (A) Gating strategy for CD45+ live cells, monocytes, B cells and γδ and αβ T cells. (B) Overview of the 50 targets analyzed with this experiment. Some of the markers can be used for phenotyping multiple immune cell lineages. (C) Representative plots for the main phenotyping markers in the B cell lineage (IgG, IgD, IgM and CD24). (D) Gating strategy to delineate invariant NKT cells, MAIT cells, CD4^+^ and CD8^+^ T cells, as well as CD4^+^ regulatory T cells (Tregs). (E) Representative plots for CD69, CD103, CD57 and PD-1 expression on non-naïve CD8^+^ cytotoxic T cells. (F) Expression pattern for CD39, CXCR3, CCR4, CD45RO and ICOS on the CD4^+^ Treg population. (G) Histogram overlays for the expression pattern of BTLA, CD27, CD28, CD38, TIGIT and KLRG1 on NK cells (grey), MAIT cells (orange), CD4^+^ Tregs (red), CD4^+^ non Tregs (purple), CD8^+^ naïve T cells (green) and CD8^+^ non-naïve T cells (blue). Dotted red lines indicate positivity cut-offs. (H) Gating strategy for NK cells and NK cell subsets based on CD56, CD161, CD16 and Nkp46. (I) Gating strategy for Basophils (CD123^+^ FcER1^+^ HLA-DR^-^), plasmacytoid DCs (CD303^+^ HLA-DR^+^), pan conventional DCs (CD11c^+^ HLA-DR^+^), and the cDC1 (CD141^+^) and cDC2 (FcER1^+^) subsets. (J) Histogram overlays for the expression pattern of CD86, CD40, CD11b, CD1c and CD163 on B cells (grey), CD8^+^ naïve T cells (green, negative control), CD14^+^ monocytes (purple), CD16^+^ monocytes (yellow), CD141^+^ cDC1s (red) and FcER1^+^ cDC2s (blue). Dotted red lines indicate positivity cut-offs. (K) Gating strategy for Lin^-^ CD2^+^ CD127^+^ innate lymphoid cells (ILCs). (L) A selection of fluorescence-minus-one (FMO) controls for the indicated markers: PD-1 and BTLA on CD8^+^ T cells, ICOS and CCR4 on Tregs, and CD86 and CD1c on Lin-cells as indicated. Dotted red lines indicate positivity cut-offs. Note that there is no or only negligible spreading error (SE) present. Additional FMO and gating controls are shown in Online Figure 9.

B cells are identified by the lineage markers CD19 and CD20 (Figure 1A). Basic B cell differentiation status can be assessed using expression of the immunoglobulin subclasses IgM, IgD, IgG and the sialoglycoprotein CD24 (Figure 1C), as well as CD27 and CD38 (11).

Pan T cells are identified by expression of CD3 (Figure 1A), followed by subsetting γδ T cells from the larger fraction of αβ T cells. MAIT cells are gated using CD161 and the invariant TCR alpha chain, TCR Vα7.2 (Figure 1D). Invariant NK-T cells can be identified using an antibody against the Vα24 and Jα18 of the TCR alpha locus. For conventional αβ T cells, after gating CD4^+^ and CD8^+^ T cells, Tregs can be identified by high expression of the IL-2 receptor alpha chain (CD25) and low expression of the IL-7 receptor (CD127) (Figure 1D). In the CD4^+^ Th cell and CD8^+^ cytotoxic T cell fraction, naïve and memory subsets can be identified by the differential expression of CD45RA, CD45RO and the chemokine receptor CCR7 (CD197) (12). Furthermore, tissue-resident memory T cells (TRMs), can be identified by the expression of CD69 and the integrin CD103 (13) (Figure 1E). CD69 also functions as a marker of recent T cell activation.

For detailed assessment of Treg function and phenotype, the ectonucleoside triphosphate diphosphohydrolase CD39 and inducible costimulator of T cells (ICOS) can be used, together with the chemokine receptors CXCR3 and CCR4 (Figure 1F) (14).

Both CD4^+^ helper T cells and CD8^+^ cytotoxic T cells can differentiate into different effector and memory lineages, but they can also enter an exhausted state during chronic infections or cancer development (15). In our panel, several phenotyping markers allow us to assess the functional state of T cells in depth: the exhaustion and activation marker Programmed Death 1 (PD-1, CD279), the senescence marker CD57, the co-inhibitory receptor BTLA (CD274), the co-receptors CD27 and CD28, the cyclic ADP ribose hydrolase CD38, TIGIT (T cell immunoreceptor with Ig and ITIM domains) and killer cell lectin-like receptor subfamily G member 1 (KLRG1). Figure 1G depicts well-defined separation for all these markers across multiple T cell subsets, and NK cells as a reference population. Furthermore, for CD4^+^ Th cells the two main effector lineages can be distinguished using the expression of the chemokine receptors CXCR3 (mostly expressed on Th1 cells) and CCR4 (mostly expressed on Th2 cells). These chemokine receptors can also be used to study different functional capacity and homing properties of CD8^+^ cytotoxic T cells and Tregs (Figure 1F).

NK cells are gated using the lineage-defining molecules CD56 (NCAM) and CD161 (KLRB1) and can be subsetted by the expression of CD16 (Figure 1H). Nkp46 functions as an additional NK cell marker that is suitable for studying tissue-derived NK cells (16).

In the myeloid cell compartment, CD14 (the LPS receptor) and CD16 (FcγRIII) are used traditionally to distinguish classical (CD14^+^ CD16^+^), intermediate (CD14^+^ CD16^dim^) and non-classical (CD14^-^CD16^+^) monocytes (17) (Figure 1A). FcER1 together with the IL-3 receptor alpha chain (CD123) are commonly used markers for basophils. Within the Lin-(CD3^-^CD19^-^CD56^-^CD14^-^CD16^-^) HLA-DR^+^ compartment, our panel identifies plasmacytoid dendritic cells (pDCs) by the expression of CD303, while classical DCs (cDCs) are marked as CD11c^+^ HLA-DR^+^. The cross-presenting cDC1 and the cDC2 subsets are identified by CD141 (also known as BDCA-3) and CD1c (BDCA-1) expression (18), respectively, with FcER1 functioning as an additional and more distinct marker for the cDC2 lineage (9) (Figure 1I). DC activation status can be assessed by the expression of the co-receptors CD40, CD86 and the Integrin alpha M (CD11b). Furthermore, CD163 allows the separation of the recently defined DC3 subset (8,19) and to phenotype monocytes (Figure 1J). Finally, ILCs can be identified by co-expression of CD127 and CD2 (Figure 1K).

A representative gating tree with all the above-mentioned subsets is shown in Figure 1, including some fluorescence-minus-one (FMO) controls: on CD8^+^ T cells for the molecules PD-1, BTLA; on CD25^+^ CD127^-^ Tregs for ICOS, CCR4; and on pan CD11c^+^ MHCII^+^ cDCs for CD86 and CD1c (Figure 1L). All of these FMOs highlight that by using systematic panel design there is negligible spreading error (SE) (20) for these populations and markers of high interest.

Our panel development strategy was based on established best practices (21–23) and multiple novel approaches. First, we utilized the similarity index (Online Figure 3), the fluorochrome brightness (Online Figure 4) and a newly developed automated algorithm for systematic fluorochrome selection. While previously described metrics such as the complexity index (24) assign an overall “score” to a given set of fluorochromes, our strategy allowed us to identify the best feasible combinations of fluorochromes to move beyond 40 colors. Second, we developed a new metric to evaluate unmixing-dependent spreading error that occurs in highly complex spectral flow cytometry panels and affects all events in the measurement (Online Figure 5, and Mage and Mair, manuscript in preparation). Finally, we utilized the instrument-specific spillover-spreading matrix (SSM) (20) and the total spread matrix (TSM) (Corselli et al, manuscript in preparation) for the optimal assignment of fluorochromes based on the biological co-expression of markers (Online Figure 6).

All steps of this panel design process, including the novel approaches, are described in detail in the online material of this manuscript, including additional staining and gating controls (Online Figure 9). Of note, this panel was developed on two full spectrum cytometers in parallel: a 7-laser instrument with a total of 186 detectors (commercially available from Sony Biotechnology as the Sony ID7000™) and a 5-laser instrument with a total of 78 detectors (commercially available from BD Biosciences as the BD FACSDiscover™ S8). The final and fully optimized panel as shown in Figure 1 was acquired on the BD FACSDiscover™ S8 (instrument configuration and setup details are listed in Online Figure 1 and 2), together with the FMO control samples. This instrument also allowed cell sorting, highlighting that 50-parameter sorting is feasible to allow very fine-grained isolation of any immune population of interest (sorting strategy and purity of the sorted populations shown in Online Figure 14). Furthermore, we established a trimmed-down panel version of 45 colors that is cross-platform compatible on the BD FACSDiscover™ S8 and Sony ID7000™ (Online Figure 12). To the best of our knowledge, this is the first report of a high-dimensional 40-color+ panel that is usable across multiple independent spectral cytometry platforms.

Manual analysis is critical to assess data quality and separation of markers in any flow cytometry panel (25), but in many cases additional insights can be gleaned from high-dimensional panels by using exploratory computational tools such as dimensionality reduction and clustering (26–28). In Online Figure 15 we show a UMAP plot with overlaid FlowSOM (29) clusters, heatmaps of the annotated clusters and histogram overlays of the raw data for some markers, highlighting the fine-grained cellular heterogeneity that can be revealed using our 50-color panel.

Overall, our data shows that this panel can serve as a widely usable and powerful immunophenotyping resource for comprehensive analysis of human immune cells in peripheral blood and non-lymphoid tissues, including human tissues of small size such as biopsies or some resected tissues (3,5).

The opportunity to reliably analyze 50 different target molecules (with the option to perform parallel cell sorting) in a high-throughput fashion is likely to enable previously impossible avenues to study the human immune system (30). Finally, development of such a comprehensive panel may enable the consistent use of a single panel suitable for multiple different studies. Together with deposition of these data into publicly accessible databases, such a consistent use would facilitate subsequent cross-study analyses with machine learning approaches such as FAUST (31,32) or other suitable computational techniques.

### Similarity to published OMIPs

The most similar OMIP to our manuscript is OMIP-069 (the first 40-color OMIP to be reported, (24)) and OMIP-044 (the first 28-color OMIP reported (33)). There is some overlap with published 28-color OMIPs focusing on T cell phenotyping (e.g. OMIP-050 and OMIP-058 (34)) and several other lower dimensional OMIPs focused on T cells, but up to date there is no OMIP that reports the use of 50 different fluorochromes allowing such in-depth phenotyping of T cells and antigen-presenting cells (APCs).

## Acknowledgements

This research was supported by the Flow Cytometry Shared Resource (RRID:SCR_022613) of the Fred Hutch/University of Washington/Seattle Children’s Cancer Consortium (P30 CA015704). We would like to thank Michele Black for her state-of-the-art flow cytometry expertise and exemplary leadership in running the Flow Cytometry Shared Resource. We thank Xiaoshan “Shirley” Shi (BD Biosciences) for assistance with equipment, and Jolene Bradford (Thermo Fisher) for support with testing reagents. This work was supported by the Emerson Collective and NIH grants R01 AI123323 and R56 DE032009 (to Martin Prlic). A.J.T. is an ISAC Marylou Scholar. P.M. is an ISAC International Innovator. F.M. is a former ISAC Marylou Scholar.

## Author Contributions

Andrew Konecny: Conceptualization; data curation; formal analysis; methodology; validation; visualization; writing - original draft; writing - review and editing.

Peter Mage: Conceptualization; formal analysis; methodology; software; validation; visualization; writing - original draft; writing - review and editing

Aaron Tyznik: Conceptualization; data curation; formal analysis; validation; project administration; resources; supervision visualization; writing - review and editing.

Martin Prlic: Conceptualization; formal analysis; validation; visualization; project administration; resources; supervision; writing - original draft; writing - review and editing.

Florian Mair: Conceptualization; data curation; formal analysis; methodology; validation; visualization; writing - original draft; writing - review and editing.

## Embedded Figure and Figure legend

## Online material

### Instrument Configuration, optimization and spectral unmixing

This panel was developed in parallel on a BD Biosciences FACSDiscover™ S8 and a Sony ID7000™. For the representative panel run shown in main Figure 1, data was acquired on a 5-laser BD FACSDiscover™ S8 Cell Sorter running BD FACSChorus™ acquisition software. The instrument contains 5 lasers with 6 spatially separated laser intercepts (the blue laser is split to support full-spectrum detection at one intercept and BD CellView™ imaging detection at a second intercept, see (1)). For the full-spectrum detection, the system utilizes a 349 nm (“UV”) laser, a 405 nm (“Violet”) laser, a 488 nm (“Blue”) laser split to two intercepts, a 561 nm (“Yellow-green”) laser, and a 637 nm (“Red”) laser. All lasers use flat-top beam-shaping optics to yield uniform illumination across the core stream. Fluorescence collection is performed via an objective lens gel-coupled to a quartz sample cuvette; the objective lens is fiber-coupled to each laser’s detector arrays.

#### General instrument description

The BD FACSDiscover™ S8 utilizes a total of 86 separate optical detectors using a combination of avalanche photodiodes (APDs), photomultiplier tubes (PMTs), and photodiodes (PDs). The 78 spectral detection channels are APDs split across 5 laser lines, along with 3 fluorescence imaging channels (PMTs) off of the 488 nm laser, 3 scatter imaging channels (2 PDs and 1 PMT) off of the 488 nm laser, and 2 scatter channels (1 PD and 1 PMT) off of the 405 nm laser (**Online Figure 1**). The optical detection paths and signal processing pathways for the 78 spectral fluorescence detectors and 3 BD CellView™ imaging fluorescence detectors are separate and independent (1). Fluorescence imaging detector data is not used for spectral unmixing, and as such the fluorescence imaging data was not used for the analysis of the data presented in this manuscript and is not discussed in further detail.

**Online Figure 1:**
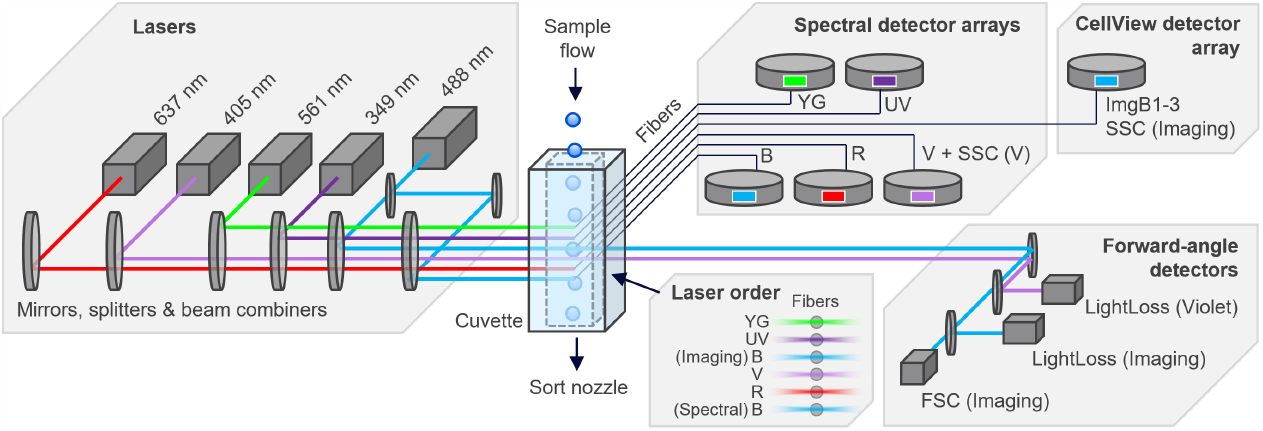
Simplified overview of the optical layout for the BD FACSDiscover™ S8. This BD FACSDiscover™ S8 was equipped with 5 spatially separate laser lines and corresponding spectral detector arrays. The data from the fluorescence imaging detector system has not been used for the data presented in this manuscript. The optical layout of the spectral detector arrays is described in more detail in Online Fig. 2 and Online Table 1.

Full-spectrum fluorescence detection (365-860nm) on the BD FACSDiscover™ S8 is achieved through a combination of bandpass filters, with the APDs arranged in a modular star-shaped pattern to minimize the number of reflections and total optical path length required. Each laser line has a dedicated and spatially separated spectral detector array, but all arrays share a common subset of detection bands depending on the total wavelength range needed (**Online Figure 2** and **Online Table 1**).

**Table 1.** embedded:

|  |  |
| --- | --- |
| **Konecny et al, OMIP-XX**<br>**Table 1. Summary table for application of OMIP XX** |  |
| **Purpose** | 50-color phenotyping of antigen-presenting cells and T cells |
| **Species** | Human |
| **Cell Types** | PBMCs and human tissue |
| **Cross-References** | OMIP-069, OMIP-044, OMIP-050, OMIP-058 and others |

**Table 2.** embedded:

| Konecny et al., OMIP-XX<br>Table 2. Reagents used for OMIP-XX |  |  |  |  |
| --- | --- | --- | --- | --- |
| Specificity | Alternative Name | Clone | Fluorochrome | Purpose |
| CD45RA | Isoform of CD45 | HI100 | Spark UV 387 | Phenotyping of T cells, naive vs memory |
| CD40 | TNFRSF5 | 5C3 | BUV395 | Phenotyping of B cells and DCs, activation |
| CD3 | Part of the TCR complex | UCHT1 | BUV496 | Lineage marker of pan T cells |
| CD56 | Neural cell adhesion molecule 1, NCAM1 | NCAM16.1 | BUV563 | Lineage marker of pan NK cells |
| CD2 | LFA-2 | S5.5 | Qdot 605 | Identification of ILCs, NK cell phenotyping |
| CD141 | BDCA-3, or Thrombomodulin | 1A4 | BUV615 | Lineage marker of cDC1s |
| Nkp46 | CD335, Natural cytotoxicity triggering | 9E2 | Qdot 625 | Phenotyping of NK cells |
| CD303 | Clec4c | V24-785 | BUV661 | Lineage marker of pDCs |
| CD86 | B7-2 | FUN-1 | BUV737 | Phenotyping of DCs and monocytes, activation |
| CD45 | Protein tyrosine phosphatase, receptor | HI30 | BUV805 | Pan-Haematopoietic marker |
| CD161 | Killer cell lectin-like receptor subfamily B 1, | DX12 | BV421 | Phenotyping of T cells and NK cells, MAIT marker |
| IgM | Immunoglobulin M | SA-DA4 | Super Bright 436 | Phenotyping of B cells and DCs |
| CD1c | N.A | L161 | Pacific Blue | Phenotyping of DCs and monocytes |
| CD28 | N.A | CD28.2 | BV480 | Phenotyping of T cells |
| CD19 | N.A | SJ25C1 | AmCyan | Lineage marker of B cells |
| CD11c | Integrin alpha X, ITGAX | S-HCL-3 | BV510 | Lineage marker for conventional DCs |
| CD45RO | Isoform of CD45 | UCHL1 | BV570 | Phenotyping of T cells, naive vs memory |
| CD197 | CCR7, chemokine receptor 7 | G043H7 | BV605 | Phenotyping of T cells, naive vs memory |
| CD69 | N.A | FN50 | BV650 | Phenotyping of T cells, tissue residency marker |
| FcER1 | high-affinity IgE receptor | AER-37 | BV711 | Lineage marker for cDC2s and basophils |
| CD103 | Integrin alpha E, ITGAE | Ber-ACT8 | BV750 | Phenotyping of T cells, tissue residency marker |
| CD194 | CCR4, chemokine receptor 4 | 1G1 | BV786 | Phenotyping of T cells, migration |
| CD25 | Interleukin-2 receptor alpha chain, IL2RA | BC96 | BB515 | Phenotyping of T cells, marker for Tregs |
| CD14 | Lipopolysaccharide receptor | M5E2 | RB545 | Lineage marker for monocytes |
| CD8 | N.A | OKT8 | NovaFluor Blue 555 | Lineage marker for CD8+ T cells |
| CD4 | N.A | SK3 | NovaFluor Blue 585 | Lineage marker for CD4+ T cells |
| CD272 | B- and T-lymphocyte attenuator, BTLA | J168-540 | RB613 | Phenotyping of T cells, activation |
| CD27 | TNFRSF7 | M-T271 | BB660 | Phenotyping of T cells, activation |
| CD11b | Integrin alpha M, ITGAM | M1/70 | PerCP | Phenotyping of APCs |
| IgG | Immunoglobulin G | G18-145 | BB700 | Phenotyping of B cells, differentiation |
| TCRgd | Gamma delta T cell receptor | B1 | RB705 | Lineage marker of gd T cells |
| CD127 | Interleukin-7 receptor subunit alpha, IL7RA | HIL7R- | RB744 | Phenotyping of T cells, Treg identification |
| TIGIT | T cell immunoreceptor with Ig and ITIM | TgMab-2 | RB780 | Phenotyping of T cells and NK cells |
| IgD | Immunoglobulin D | W18340F | PerCP-Fire 806 | Phenotyping of B cells, differentiation |
| CD57 | Human natural killer-1, HNK-1 | NK-1 | PE | Phenotyping of T cells and NK cells |
| CD20 | N.A | 2H7 | Spark YG 593 | Lineage marker of B cells |
| CD24 | Signal transducer CD24 | SN3 | PE-Alexa Fluor 610 | Phenotyping of B cells, differentiation |
| CD183 | CXCR3, CX chemokine receptor 3 | G025H7 | PE-Fire 640 | Phenotyping of T cells, migration |
| CD123 | Interleukin-3 receptor, IL3RA | 9F5 | PE-Cy5 | Lineage marker of Basophils and pDCs |
| CD278 | Inducible T-cell costimulator, ICOS | ISA-3 | PE-Cy5.5 | Phenotyping of T cells, activation |
| CD16 | Fc gamma receptor, FcγRIII | 3G8 | PE-Alexa Fluor 700 | Lineage marker of monocytes, phenotyping of NK |
| CD279 | PD-1, Programmed Death 1 | EH12.1 | PE-Cy7 | Phenotyping of T cells, activation and exhaustion |
| HLA-DR | MHC class II | L243 | PE-Fire 810 | MHC class II |
| Va24-JA18 | V alpha 24 – J alpha 18 TCR chain | 6B11 | APC | Marker for invariant NKT cells |
| Va7.2 | V alpha 7.2 TCR chain | 3C10 | Alexa Fluor 647 | Marker for MAIT cells |
| KLRG1 | Killer cell lectin-like receptor subfamily G | SA231A2 | Spark NIR 685 | Phenotyping of T cells and NK cells, activation |
| CD39 | Ectonucleoside triphosphate | A1 | R718 | Phenotyping of T cells and NK cells, activation |
| Live/Dead | N.A | Amine | Zombie-NIR | Live/Dead cell discrimination |
| CD163 | Scavenger receptor for hemoglobin | GHI/61 | APC-Cy7 | DC phenotyping marker, marker for DC3s |
| CD38 | cyclic ADP ribose hydrolase | HB-7 | APC-Fire 810 | Phenotyping of T cells and NK cells, activation |

**Online Figure 2:**
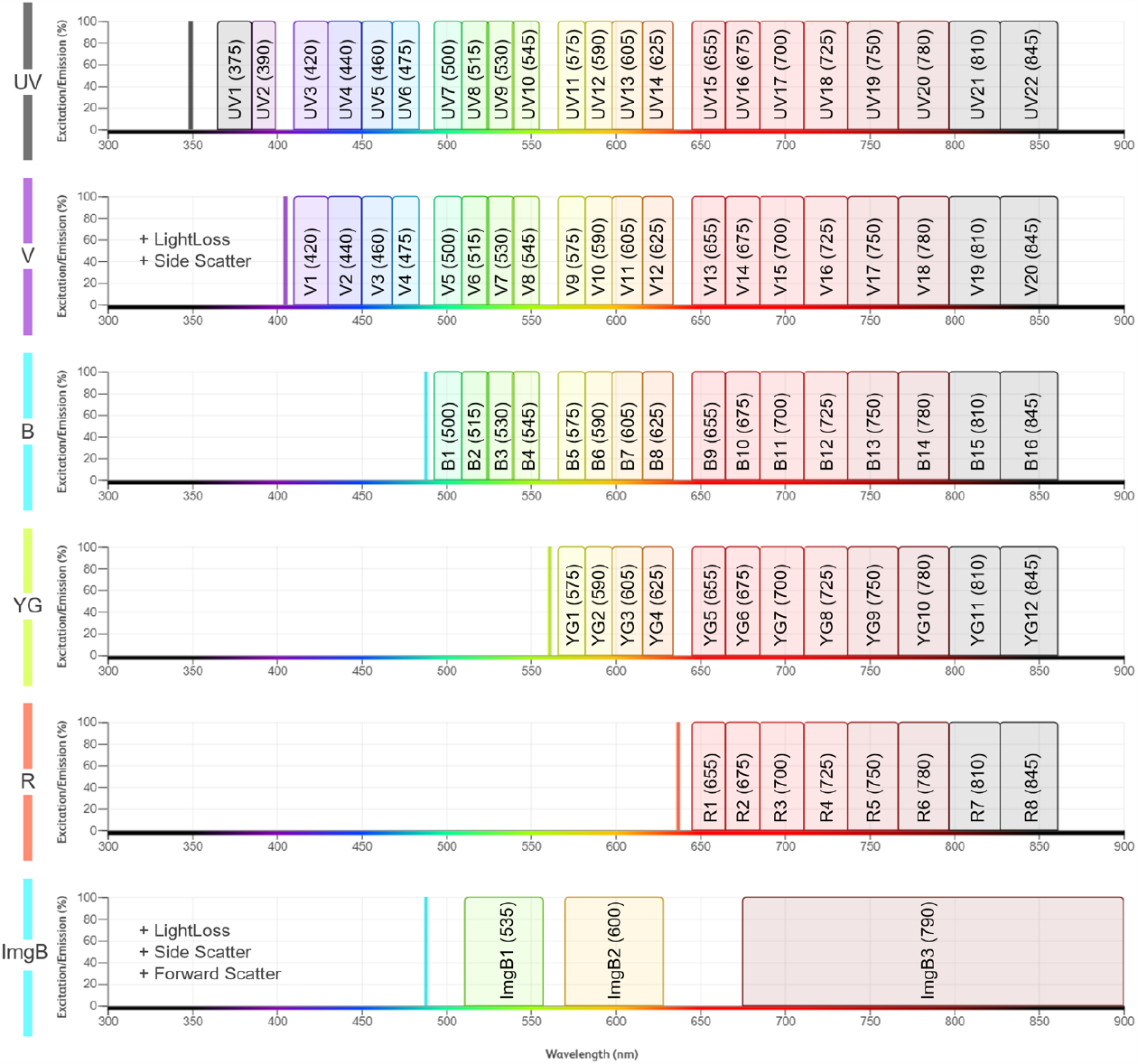
Graphical representation of the detector layout on the BD FACSDiscover™ S8. This overview shows how common subsets of detector bands are used across each spectral detector array. Vertical lines represent the wavelength of the 5 laser lines, with a gap in the detector arrays preventing the collection of any scattered laser light. The precise wavelength cut-offs for each detector band are listed in Online Table 1. Data from the imaging module was not utilized in this manuscript.

The BD FACSDiscover™ S8 reports for each event on each detector the following parameters: pulse area (-A), pulse height (-H), pulse width (-W), and pulse time-to-peak (-T). Spectral unmixing is performed on pulse area measurements from the system’s 78 spectral detector bands as described in more detail below.

**Online Table 1:** Detailed list of the detector bands used in the spectral detector arrays of the BD FACSDiscover™ S8. The name and approximate central wavelength for each spectral detector is listed for all five laser lines (first five columns). The last two columns list the effective central wavelength (CWL) and the effective bandwidth (BW) of each detector. The effective CWL and effective BW for a given detector band are determined by the specific arrangement of bandpass filters in the optical path leading to that detector.

| Ultraviolet<br>349 nm / 30 mW | Violet<br>405 nm / 50 mW | Blue<br>488 nm / 50 mW | Yellow-Green<br>561 nm / 50 mW | Red<br>637 nm / 100 mW | Effective<br>CWL (nm) | Effective<br>BW (nm) |
| --- | --- | --- | --- | --- | --- | --- |
| UV1 (375) | - | - | - | - | 375 | 20 |
| UV2 (390) | - | - | - | - | 391 | 12.5 |
| UV3 (420) | V1 (420) | - | - | - | 420 | 20 |
| UV4 (440) | V2 (440) | - | - | - | 440 | 20 |
| UV5 (460) | V3 (460) | - | - | - | 459 | 18 |
| UV6 (475) | V4 (475) | - | - | - | 476 | 16 |
| UV7 (500) | V5 (500) | B1 (500) | - | - | 501 | 16 |
| UV8 (515) | V6 (515) | B2 (515) | - | - | 517 | 16 |
| UV9 (530) | V7 (530) | B3 (530) | - | - | 532 | 16 |
| UV10 (545) | V8 (545) | B4 (545) | - | - | 547 | 16 |
| UV11 (575) | V9 (575) | B5 (575) | YG1 (575) | - | 575 | 14 |
| UV12 (590) | V10 (590) | B6 (590) | YG2 (590) | - | 590 | 16 |
| UV13 (605) | V11 (605) | B7 (605) | YG3 (605) | - | 607 | 18 |
| UV14 (625) | V12 (625) | B8 (625) | YG4 (625) | - | 623 | 14 |
| UV15 (655) | V13 (655) | B9 (655) | YG5 (655) | R1 (655) | 655 | 20 |
| UV16 (675) | V14 (675) | B10 (675) | YG6 (675) | R2 (675) | 675 | 20 |
| UV17 (700) | V15 (700) | B11 (700) | YG7 (700) | R3 (700) | 698 | 26 |
| UV18 (725) | V16 (725) | B12 (725) | YG8 (725) | R4 (725) | 724 | 26 |
| UV19 (750) | V17 (750) | B13 (750) | YG9 (750) | R5 (750) | 752 | 30 |
| UV20 (780) | V18 (780) | B14 (780) | YG10 (780) | R6 (780) | 782 | 30 |
| UV21 (810) | V19 (810) | B15 (810) | YG11 (810) | R7 (810) | 812 | 30 |
| UV22 (845) | V20 (845) | B16 (845) | YG12 (845) | R8 (845) | 844 | 34 |

#### Spectral unmixing

The BD FACSDiscover™ S8 utilizes a proprietary spectral unmixing algorithm called SpectralFX™ System-Aware unmixing. This algorithm is designed to manage spreading error (SE) in unmixed data by adapting to instrument performance and sample conditions in real-time. The unmixing algorithm is sort-compatible. Offline data analysis in FlowJo 10.10 (BD Biosciences) may be performed using either SpectralFX™ unmixing or using traditional ordinary-least-squares (OLS) unmixing, both of which are available in the exported FCS files. The data shown in Figure 1 and all Online figures with BD FACSDiscover™ S8 data has been unmixed using SpectralFX™ unmixing, unless stated otherwise.

**Online Figure 3:**
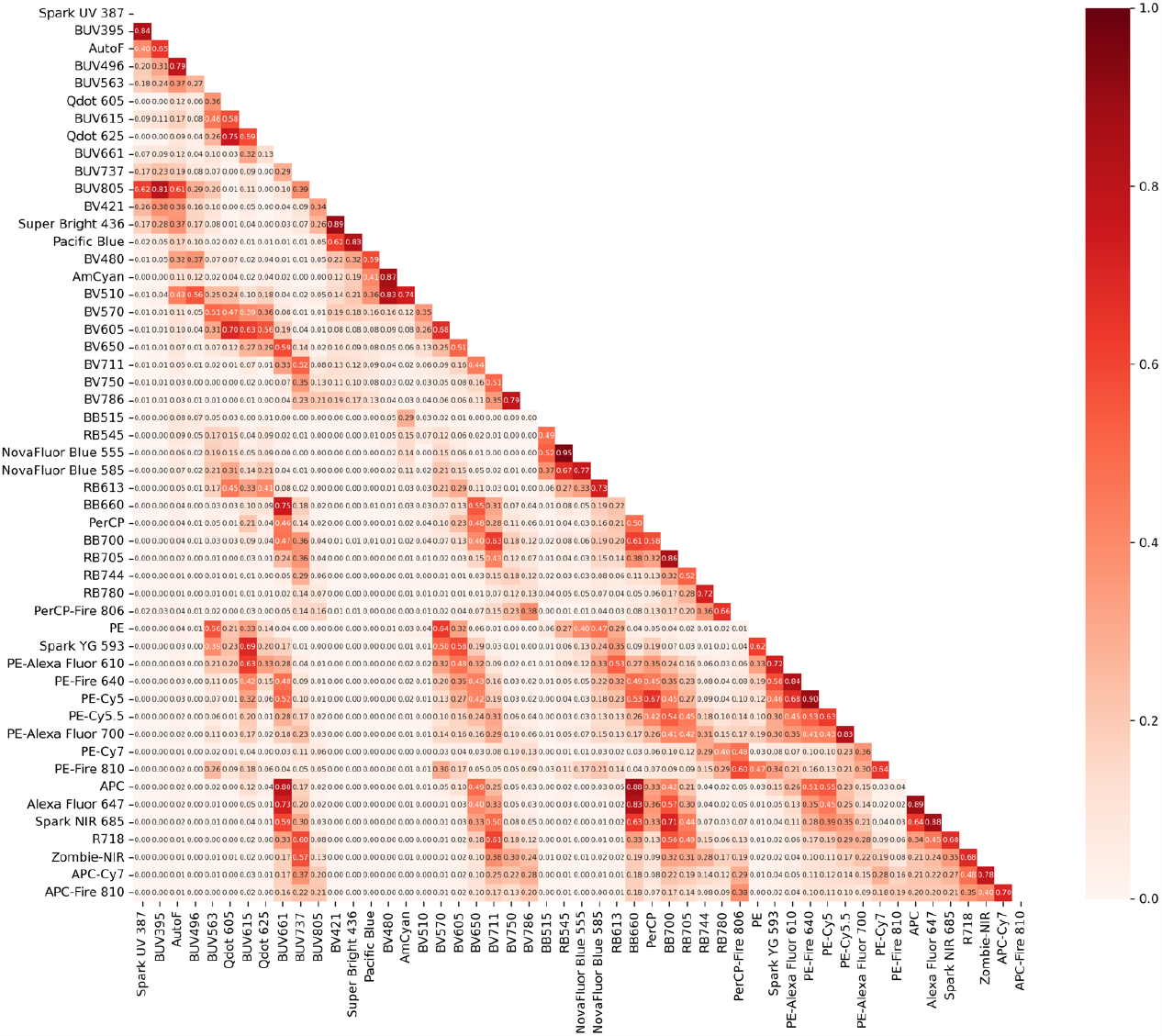
Similarity index for the final fluorochrome selection on the BD FACSDiscover S8. PBMCs were stained with anti-human CD4 antibodies conjugated to the indicated fluorochromes and acquired on a BD FACSDiscover™ S8 at standard settings. Fluorochrome similarity index was calculated as the cosine of the angle of the fluorochrome vector across the entire detector space.

**Online Figure 4:**
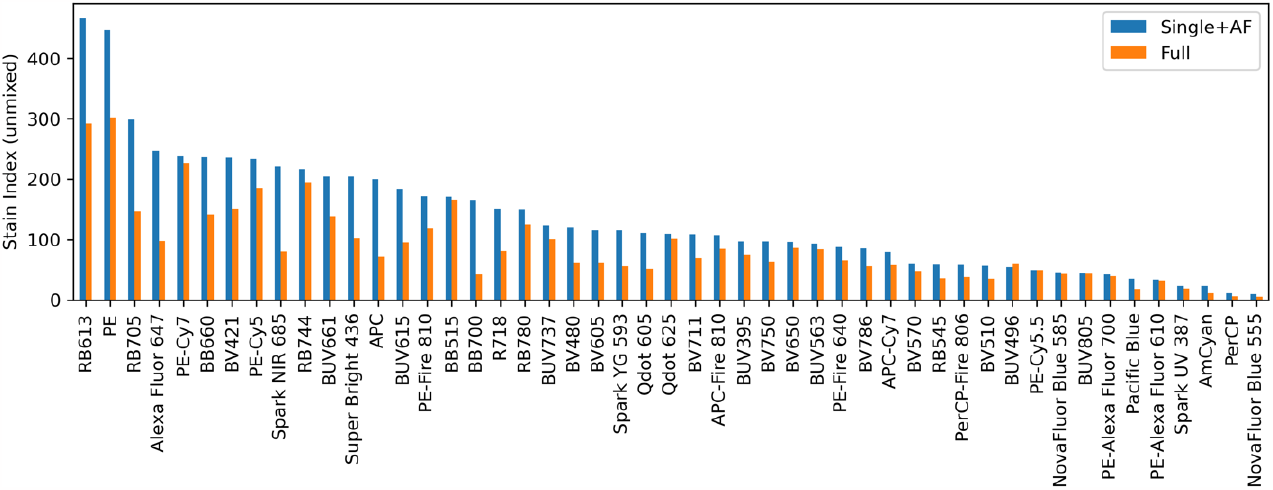
Fluorochrome resolution chart on the BD FACSDiscover S8. PBMCs were stained with anti-CD4 antibodies conjugated to the indicated fluorochromes, and acquired on a BD FACSDiscover™ S8 at standard settings. The individual single-stains were unmixed either by themselves (with Autofluorescence, blue bars) or in the context of the full 50-color unmixing matrix (orange bars). Fluorochromes are ordered based on descending stain index (SI) using the single-color unmixing. Stain index was calculated using the standard formula 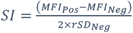.

#### Instrument setup and optimization

The BD FACSDiscover™ S8 uses a combination of an on-board broad-spectrum LED pulser (2) and a broad-spectrum, single-peak, hard-dyed setup bead (BD FACSDiscover™ Setup Bead) to perform detector setup and quality control. The LED pulser is used to independently measure sensitivity on each detector and automatically calibrate detector voltage-gain response curves for all detectors on the system. Consequently, gain control within BD FACSChorus™ software is implemented in calibrated decibel (dB) units using the formula *Gain*_*dB*_ = 20 × log_10_(*Gain*_*linear*_), such that an increase of 20 dB corresponds to a 10-fold increase in linear detector gain and a corresponding 10-fold increase in measured signal. Default APD gain settings are established during the instrument’s daily setup routine to achieve consistent signal from the FACSDiscover™ Setup Bead at a pre-defined target -Area MFI in each detector channel.

Default APD gains were chosen by the manufacturer to balance detector sensitivity (defined as the dimmest signal resolvable from noise) and detector dynamic range (defined as the brightest on-scale signal). This was achieved by evaluating detector electronic and optical noise at a range of gain settings, as well as stain index and MFI of PBMCs stained with a wide range of anti-CD4 conjugated fluorochromes spanning the full range of excitation laser wavelengths and emission detector wavelengths on the system (data not shown).

All BD FACSDiscover™ S8 data presented in this manuscript was acquired at the default APD gain settings (as optimized by the manufacturer), and the populations of interest in all samples were determined to be fully on-scale (defined as having -Height MFI <6.5*10^4^) in all spectral detector channels at those default gains. Notably, all BD FACSDiscover™ S8 instruments use calibrated detector gains and target the same setup bead MFIs by default, suggesting that detector gain adjustment and re-optimization may not be required for reproduction of the panel results presented in this manuscript on additional BD FACSDiscover™ S8 instruments.

### Strategy for Panel development

To develop this panel in a systematic way, we followed a three-step process. First, we evaluated the available fluorochrome landscape using a combination of new and existing metrics with the goal of describing the likely performance of fluorochromes in the context of a full 50-color panel. Second, we used these metrics to select a suitable palette of 50 fluorochromes through a combination of manual iterative selection and a novel automated fluorochrome selection algorithm. Finally, we combined our fluorochrome performance metrics with an understanding of the major biological expression patterns in our panel to assign the fluorochromes in our 50-color palette to individual markers in our panel.

Throughout the panel development process, we periodically tested preliminary versions and subsets of the panel, iteratively refining and updating the panel design based on full-panel performance. Each step in the process is described in more detail below.

#### Metrics for fluorochrome evaluation

For systematic fluorochrome evaluation, we used three categories of performance metrics. The first category includes metrics that describe fluorochrome performance in isolation or in pairs, but are independent of a panel’s full color palette (the full set of fluorochromes in the panel).

The second category contains metrics that describe fluorochrome performance in the context of spectral unmixing within a full panel’s color palette, but are agnostic to marker coexpression patterns in the biological sample of interest. The third category includes metrics that indicate the performance of a fluorochrome in the context of a full panel with potential marker coexpression.

#### Palette-independent fluorochrome metrics

First, we tested a large set of fluorochromes (more than 100, data not shown) as anti-CD4 conjugates on both the Sony ID7000™ and the BD FACSDiscover™ S8 to assess the spectral similarity and exclude fluorochromes that were either too similar for meaningful spectral unmixing or too dim. A heatmap representing the similarity index for the final selection of fluorochromes on the BD FACSDiscover™ S8 is shown in **Online Figure 3**. The highest similarity index at 0.95 was the fluorochrome pair RealBlue 545 and NovaFluor Blue 555, which we assigned to mutually exclusive cell lineage markers (CD14 on monocytes, and CD8 on T cells, respectively). All other similarity indices were less than 0.90.

Furthermore, we calculated the relative fluorochrome resolution (determined by both brightness and unstained population variance, or “negative spread”) for this selection of fluorochromes (**Online Figure 4**). Of note, while fluorochrome brightness is panel-independent, relative fluorochrome resolution is indirectly affected by the complexity of the unmixing matrix (single-color unmixing vs 50-color unmixing), as explained in more detail in the next section.

#### Palette-dependent fluorochrome metrics

Second, we sought to understand how specific combinations of fluorochromes would perform in the context of full-panel unmixing. The condition number (CN) of a panel’s spectral matrix, also called the panel’s complexity index, has previously been reported to predict panels with potentially problematic combinations of fluorochromes (3). Although useful as a single summary metric of panel usability, the CN does not indicate which fluorochromes are implicated in unmixing issues, nor does it quantify the severity of potential spreading issues within the panel. As such, the CN is not helpful in identifying particular areas of concern for a given panel.

**Online Figure 5:**
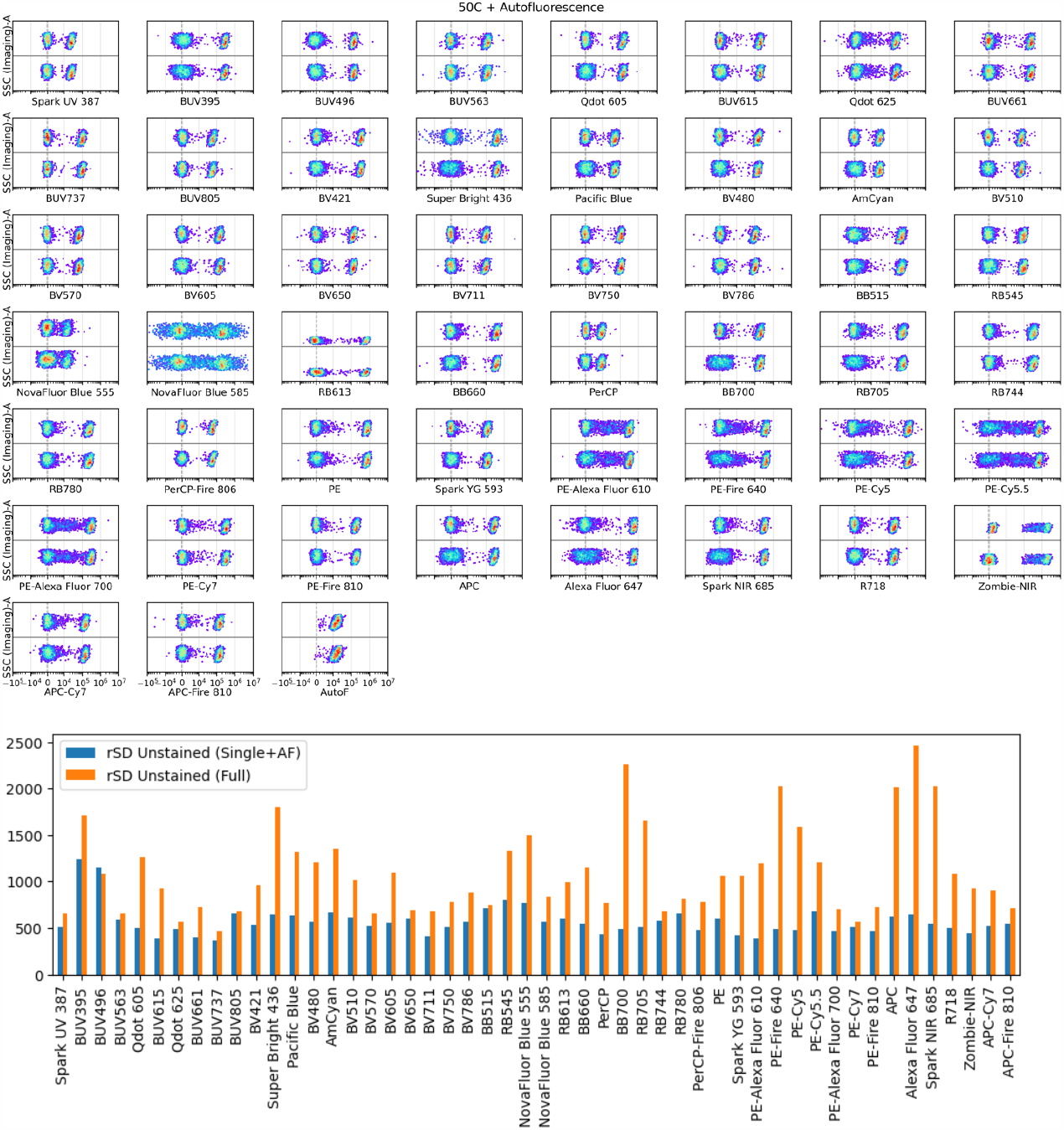
The effect of unmixing spreading error on the rSD of the negative population (I.e. all cells in the sample). The top panel shows the individual single-stains unmixed either by themselves (with Autofluorescence, top of each plot) or in the context of the full 50-color unmixing matrix (bottom of each plot). The lower panel shows the quantification of the rSD for each of these single-stains under the two unmixing conditions (single with Autofluorescence: blue bars vs full 50-color unmixing: orange bars). Note how some fluorochromes (e.g. PE-Cy7 or BB515) are not affected by spectral unmixing error, while some fluorochromes (e.g. BB700) show up to a 5-fold increase in rSD.

Therefore, we also deployed a new metric that we termed “unmixing spreading error”. This metric reveals how the complexity of the spectral unmixing matrix affects the increase in noise for all measured cells on a per-fluorochrome-basis, irrespective of the presence of positively stained cells. This effect is particularly evident for panels in the 35-color+ space and, in the example of our final 50-color panel, can lead to a 5-fold loss of resolution for certain fluorochromes due to increased variance in the negative cell population (Mage and Mair, manuscript in preparation).

**Online Figure 5** (top panel) shows for each anti-CD4 single stain the observed staining pattern when unmixed in the context of single-color unmixing with Autofluorescence (AF) (top subplots) relative to unmixing of the same single-stained sample in the context of the 50-color unmixing matrix (bottom subplots). The bar graph (**Online Figure 5**, bottom panel) shows how the rSD of the negative population for the respective fluorochrome changes in each of these conditions, which allowed us to pinpoint the fluorochromes that suffer from the largest increase in variance (e.g. AF647). This increase in variance of the negative cells (i.e. “unmixing spreading error”) directly impacts the relative separation of these fluorochromes, as can be seen by the reduction in stain index when unmixing in the context of the 50-color unmixing matrix (**Online Figure 4**).

We used the conclusions from this analysis to identify unusable combinations of fluorochromes in the context of potential full panel designs, and eventually to assign fluorochromes in the final panel that would be affected by unmixing spreading error either to lineage markers or to targets that would produce a sufficiently bright signal.

Two examples that are particularly affected by unmixing spreading error are AF647 and BB700. AF647 was assigned to Vα7.2, a TCR chain expressed primarily on MAIT cells (and other CD8^+^ T cells), which produces a bi-modal signal bright enough for good separation. BB700 was assigned to IgG, which also produces a bi-modal signal on B cells that is bright enough for good separation. PE-Fire 640 and Spark-NIR 685 only had limited reagent availability, which is why these fluorochromes had to be assigned to phenotyping markers, despite being affected by significant unmixing spreading error (discussed in more detail below in the section “marker assignment”).

#### Coexpression-dependent fluorochrome metrics

Third, we calculated the spillover spreading matrix (SSM) (**Online Figure 6A**) (4) and the total spread matrix (TSM) (**Online Figure 6B**) (Corselli et al., manuscript in preparation) for the set of dyes that we used for the final panel development. The SSM formed the basis for minimizing potential spreading error issues that result from co-expression of markers on the same cell type, as has been described in several best-practice papers and previous OMIPs. Of note, the similarity index does not replace the use of the SSM or TSM for marker assignment, as the relative similarity index does not reliably correlate with the extent of spreading error (Mair and Mage, unpublished data).

#### Fluorochrome palette selection process

Using these fluorochrome performance metrics, we evaluated potential combinations of fluorochromes given fluorochrome reagent availability for the biological markers in our panel. The biological logic for our initial assignments is described below. First, a manual iterative process was used to identify a feasible palette of approximately 40 colors with acceptable performance based on the metrics described above. Beyond this point, the huge number of possible permutations for expanding to the final 50 needed colors made this manual iterative process intractable, and the condition number (i.e. the complexity index) as metric alone is insufficient (**Online Figure 7A-B)**.

**Online Figure 6:**
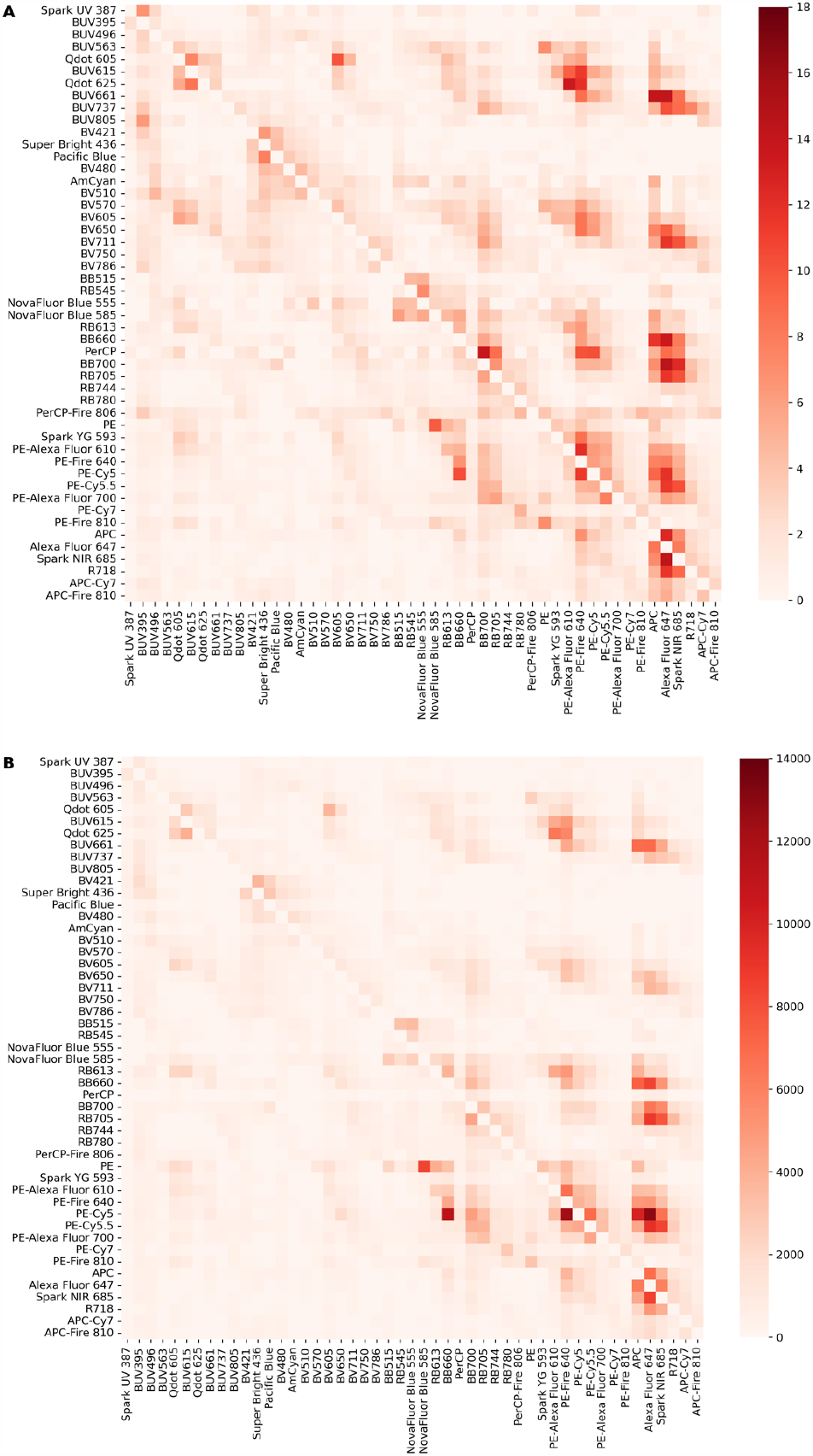
Spillover spreading matrix (SSM) and total spread matrix (TSM) for the final selection of 50 fluorochromes on the BD FACSDiscover™ S8. The spillover spreading matrix (SSM) and total spread matrix (TSM) were calculated using the same set of anti-CD4 single stains as described above. (A) Spillover spreading matrix. The fluorochrome pair with the highest SE was BB700 into Alexa Fluor 647 (SE = 14.42). Of note, SE from Alexa Fluor 647 into BB700 is negligible. (B) Total spread matrix. The TSM is calculated similarly to the SSM, except the values in the TSM are not normalized to the MFI of the positively-expressed fluorochrome, thereby revealing fluorochromes that cause significant SE as a result of both spectral overlap and brightness. The fluorochrome pair with the highest total spread error (TSE) was PE-Cy5 into Alexa Fluor 647 (TSE=12,860). The SSM and TSM were calculated based on single-stain data unmixed using the full 50C matrix (including autofluorescence). Live/dead stain (Zombie-NIR) and autofluorescence were excluded from the final SSM and TSM for clarity.

To solve this problem, we developed an automated algorithm (implemented in the Python programming language) that uses single-stain spectral signatures to identify usable 50-color combinations of fluorochromes. This algorithm finds combinations of fluorochromes that minimize metrics related to spreading error. Although the algorithm can be used to generate feasible color palettes *de novo*, we used our manually-selected ∼40-color palette (data not shown) as a starting point for generating the full 50-color panel. After constraining the fluorochrome options based on reagent availability for our markers of interest, we used the algorithm to produce a variety of possible 50-color combinations with nearly equivalent CN. We then used unmixing spreading error analysis to identify the panel options with the least impact from unmixing-dependent spread (example shown in **Online Figure 7A-B**).

This optimization process led us to a final 50-color panel and 45-color trimmed-down panel (described below) that both have substantially lower complexity than previously reported 40-color panels (**Online Figure 7C**) (3,5,6). Our final panel has a condition number of 40.2, while the 40 fluorochromes selected in OMIP-069 result in a value of 57 (on the BD FACSDiscover™ S8).

A concise visual overview for the spectra of the fluorochromes selected in the final panel is shown in **Online Figure 8**. We plotted normalized spectral signatures as line plots per primary laser excitation group (top panel) to facilitate identification of the peak detectors for each fluorochrome, and we visualized the same data as a heatmap (bottom panel) to help identify fluorochromes that show significant cross-laser excitation (e.g. Qdots, PerCP).

#### Marker assignment to fluorochromes

In terms of biology, the overarching goal of this panel was to analyze as many markers as feasible to assess the functional state of T cells and antigen-presenting cells (APCs) in diseased and healthy human blood and tissue samples. While we aimed to have a broadly applicable panel, we prioritized markers that have a known relevance in the context of T cell exhaustion, T cell migration, tissue residence and regulatory T cell differentiation. Among these high-priority markers were the co-inhibitory molecules PD-1, ICOS, TIGIT, the chemokine receptors CXCR3, CCR4 and the tissue-residency markers CD69 and CD103. Furthermore, it was a requirement to evaluate CD40 and CD86 on all known APC populations: cDC1s, cDC2s, the more recently described cDC3 subset, pDCs, classical and non-classical monocytes (in blood) and also macrophage populations (in tissues).

In parallel we aimed to capture a snapshot and basic subset distribution of the remaining main immune subsets: NK cells, B cells and innate lymphoid cells. To make the panel applicable not only to PBMCs, but also to human tissue samples, the inclusion of a sufficiently bright CD45 was mandatory. Based on the SSM and prior experience (5), CD45 was assigned to BUV805, a relatively dim dye that is not causing any issues with spreading error (SE) for all other fluorochromes. For the assignment of the remaining target markers, the following rationales were used:

**Online Figure 7:**
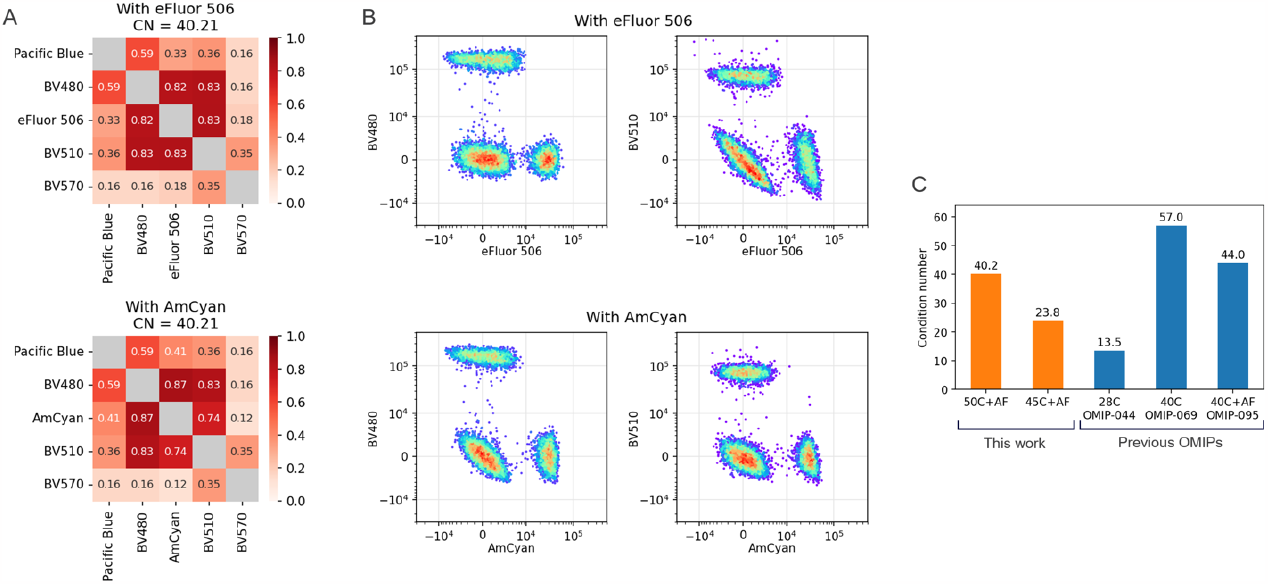
Refinement of optimal color selections through analysis of unmixing spreading error and condition number. (A) Spectral similarity indices and spectral matrix condition numbers (CN) as measured on the BD FACSDiscover™ S8 for a subset of violet-excited fluorochromes in two nearly identical versions of our 50-color panel, using either eFluor 506 (top) or AmCyan as in the final panel (bottom). Both versions have identical CN values and comparable similarity indices, with the closest spectral neighbors for both dyes being BV480 and BV510 (having similarities of 0.82 and 0.83 respectively for eFluor 506, or 0.87 and 0.74 for AmCyan). (B) Analysis of unmixing spreading error (shown here with overlaid plots of unmixed anti-CD4 single stain samples) in the context of both full panels reveals substantially worse unmixed spread and resulting loss of resolution for eFluor 506 compared to AmCyan. Based on this analysis, we chose to proceed with AmCyan in the final panel. (C) Comparison of CN for the final 50-color and 45-color panels reported in this manuscript (orange) and three previously published OMIPs (blue). CN calculations were based on spectral signatures measured on the BD FACSDiscover™ S8. Spectral signature data for Live/Dead Fixability Blue (Thermo Fisher) was not available, so calculations were performed using the spectrum of the nearly identical BD Fixable Viability Stain 440UV instead.

- CD3 was considered the key lineage marker to identify pan T cells. Based on prior experience and the SSM, BUV496 was assigned for CD3. Similarly to BUV805, this fluorochrome does not generate any SE for all other fluorochromes (rowsum less than 20), thus maintaining best possible resolution for the assignment of downstream T cell phenotyping markers.
- CD11c as well as HLA-DR were considered the key lineage markers for dendritic cells. Initial tests included CD11c-AF700 and HLA-DR-APC-Cy7. However, these widely used lineage markers were also available on more recently released fluorochromes, and thus we tested HLA-DR PE-Fire 810, which resulted in a very bright signal, and causing SE in relatively few other fluorochromes. The affected fluorochromes (PE and PE-Cy7) were assigned to T cell lineage markers, since HLA-DR is only expressed at lower levels on T cells. For CD11c, we decided to utilize BV510, a relatively dim fluorochrome that only generates significant SE for BUV496 (assigned to CD3) and the neighboring fluorochromes on the violet detector array.
- After making choices for these key lineage markers, we focused on reasonable assignments for the fluorochrome-pairs that showed very high spreading error based on the SSM: BUV661 and APC/AF647, Qdot 625 and PE-AF610/BUV615, Spark NIR 685 and AF647, and the pair AF647 and APC itself. Initial tests centered on Qdot 625, as the availability of target antigens for this fluorochrome was very limited. In the end we assigned Qdot 625 as a Streptavidin-conjugate to Nkp46, a marker of NK cells, with BUV615 assigned to the mutually exclusive CD141 (cDC1 lineage) and PE-AF610 assigned to CD24, which is primarily expressed on B cells and some myeloid cells.

**Online Figure 8:**
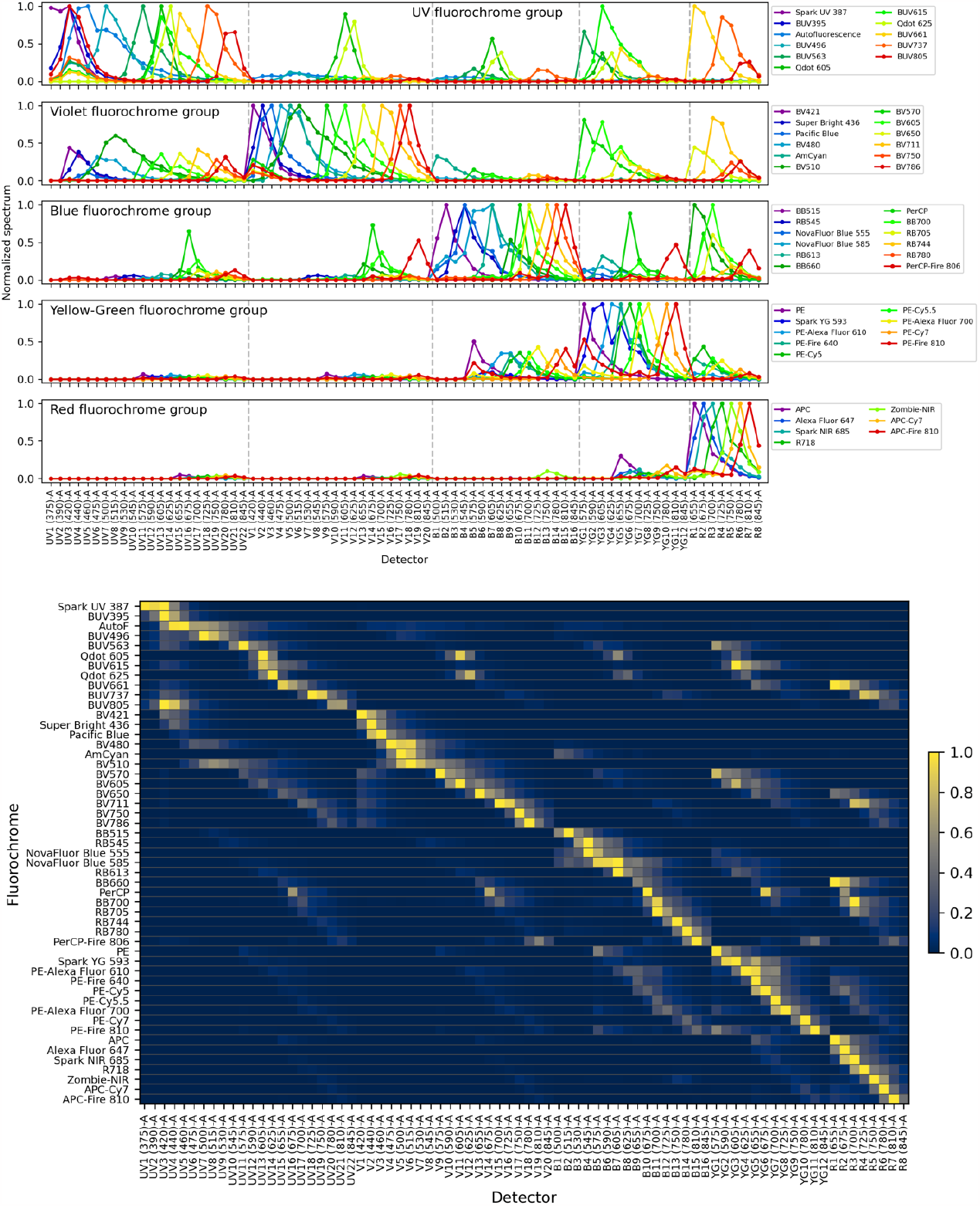
Spectral signatures for the final fluorochrome selection on the BD FACSDiscover™ S8. PBMCs were stained with anti-human CD4 antibodies conjugated to the indicated fluorochromes and acquired on a BD FACSDiscover™ S8 at standard settings. Fluorochrome spectral signatures (normalized emission spectra) are shown as line plots grouped by primary laser (top) and as a heatmap (bottom).

BUV661 was tested and assigned to CD303, a unique marker for plasmacytoid DCs (pDCs), with APC and AF647 reserved for mutually exclusive T cell lineage markers. Of note, APC as well as AF647 both are affected significantly by unmixing spreading error (see **Online Figure 4)**. However, after testing various targets we concluded that APC on Va7.2 (MAIT cells) and AF647 for Va24Ja18 (NKT cells) delivered sufficient resolution for either population. Finally, the assignment for Spark NIR 685 required multiple rounds of testing as this fluorochrome is relatively dim and also suffered a significant loss of resolution from unmixing spreading error (Online Figure 4). KLRG1 (expressed by MAIT cells and some T cells) showed high enough expression to be assigned to Spark NIR 685.

- Super Bright 436 and BV421 based on their high similarity index (0.89) were initially considered a troublesome pair, but the actual SE generated for this pair is relatively low. Nevertheless, we assigned these fluorochromes to mutually exclusive antigens: Super Bright 436 to IgM, a marker of naïve B cells, and BV421 to CD161, which is expressed on MAIT cells, NK cells and some subpopulations of T cells.
- Similarly, BB700 and RB705 with a similarity index of 0.86 were considered a problematic pair, but also here the actual SE was relatively low. BB700 was assigned to IgG, a subsetting marker for B cells, and RB705 as a very bright fluorochrome was utilized for TCRgd, a lineage marker of γδ T cells.
- Another problematic triad was PE-Cy5, PE-AF700 and PE-Fire 640, which showed high similarity indices and also a relatively high degree of SE, particularly PE-Cy5 into PE-Fire 640. Furthermore, availability of antibody-conjugates for these two dyes was very limited. After trying out different reagents, we settled on CD16-PE-AF700 (expressed on monocytes and NK cells) and CXCR3-PE-Fire 640 (a chemokine receptor only found on T cells). Of note, PE-Fire 640 suffered from significant unmixing spreading error, making it less ideal for a chemokine receptor, but limited reagent availability forced us to compromise.

The third of this fluorochrome triad, PE-Cy5, was assigned to CD123, which is the IL-3 receptor and expressed primarily on the distinct lineages of basophils and pDCs.

- In general, part of the fluorochrome selection process was limited by the availability of useful fluorochrome-antibody conjugates for more recently released dyes. For example, we utilized NovaFluor Blue 555 and NovaFluor Blue 585 for CD8 and CD4 respectively. Both of these dyes are very dim, not available with a lot of targets, and also receive some degree of spreading error (e.g. NovaFluor Blue 585 from PE). However, resolution in the full panel was considered sufficient to separate CD4 and CD8 T cells.
- Similarly, since PE-AF610 availability was very limited, we chose it for CD24 to analyze B cell subsets. The chosen reagent delivered good separation.
- For the most important functional phenotyping markers within the T cell and APC lineage, we followed the approach of minimizing spreading error received by a fluorochrome (i.e. choosing row-sums in the SSM that are minimal or within a low range) together with preferably bright fluorochromes. Of high importance to us were PD-1, TIGIT as well as CD39 for T cells; CD86 and CD40 for APCs.

Balancing fluorochrome brightness with minimal row-sums in the SSM led to a selection of about a dozen fluorochromes, of which we chose the following final selection: PD-1-PE-Cy7 (almost no SE received at all, very bright fluorochrome), TIGIT-RB780 (no SE except from PE-Cy7) and CD39-R718 (SE only from BUV737 and BV711, both of which were later assigned to myeloid cell markers). CD86-BUV737 and CD40-BUV395 (both of which only have low SE received).

CD69 and CD103 were assigned based on reagents successfully used in previous high-parameter panels (7): CD69-BV650 and CD103-BV750, both of which delivered good resolution for these two tissue-residency markers for T cells.

- BTLA had been used in a previous panel (7) conjugated to BB630. We switched this to a newer dye format that shows more specific excitation from the blue laser, RB613. Initial titrations and testing showed great separation and very high staining index for RB613 and thus we settled on BTLA-RB613.
- The T cell markers CD27 and CD28 had previously been used as BB660 and BV480, respectively. Tests with early panel iterations showed that both markers delivered the same resolution as known from lower-dimensional panels, and based on the SSM both dyes remained either free from SE (BV480) or collected SE only from fluorochromes assigned to mutually exclusive cell lineages (for BB660: PE-Cy5 assigned to CD123).
- Some fluorochromes were selected based on their beneficial profile in our initial assessment of spectral similarity and spillover characteristics, but had only very limited target antigens commercially available. These fluorochromes included APC-Fire 810, PerCP-Fire 806 and Qdot 605. Based on published work (3) we tested CD38-APC-Fire 810, which is broadly expressed on multiple cell lineages as an activation or phenotyping marker, and delivered good resolution. For PerCP-Fire 806 we decided to use IgD, expressed only on B cells. Qdot 605 was available as anti-CD2, which is expressed on T cells, ILCs, and cDC2s.
- CD127 was initially used based on prior experience as a BV786-conjugate, with intermediate performance. Based on conjugate-availability we switched to CD127-RB744 (one of the top 10 fluorochromes in terms of brightness, relatively low unmixing spreading erro), which performed very well and was selected as the final choice for this marker.
- CD25 was considered high priority as the primary marker delineating regulatory T cells (Tregs) from the remainder of T cells. Based on the fluorochrome brightness and low SE received (low column sums in the SSM) we tested BV421 and BB515, both of which performed well. For the final panel we settled on CD25-BB515.
- CD161 was assigned to BV421, delivering great resolution.
- ICOS (CD278): This was considered a key functional molecule on the Treg population and as such we tested multiple different options: Initially, ICOS-BUV737 was selected based on the low SSM column value and prior experience. While this conjugate performed well, we also tested ICOS-PE-Cy5.5, which is a fluorochrome with limited reagent availability. Despite the fact that PE-Cy5.5 sits in a relatively crowded area of the spectrum, unmixing-spreading error is not very pronounced and overall SE collected from the fluorochromes assigned to the T cell lineage was minimal. Together with the respective FMO-control we concluded that ICOS-PECy5.5 performed satisfactory.
- CD45RA and CD45RO were assigned to Spark UV 387 and BV570, respectively. CD45RA performance was sub-optimal because of Spark UV 387 being one of the three dimmest fluorochromes available, but biological resolution together with CCR7 and CD45RO was considered sufficient.
- For the chemokine receptor CCR7, we tested multiple conjugates: BV510, BUV661, PE, PE-CF594 and BV605. All of these worked comparably well, but of the remaining fluorochrome slots available BV605 was chosen.
- CD2, a molecule broadly expressed on T cells and used as a marker for innate lymphoid cells (ILCs) was one of the few targets available on Qdot 605 and performed well.
- For the PE and Spark-YG593 pair we initially expected high spreading error because of the spectral similarity of these fluorochromes, which after systematic analysis turned out to be much lower than anticipated. Nevertheless, we assigned these to mutually exclusive cell lineages and relatively bright targets: CD57 (expressed on some cytotoxic T cells) to PE and Spark-YG593 to CD19 (the main lineage marker of B cells).
- CD20, another lineage defining marker for B cells was assigned to Amcyan, a fluorochrome that did not cause any SE for the fluorochrome-marker combinations selected for B cell phenotyping (CD24, CD38, IgG, IgD, IgM).
- For unambiguous identification of different dendritic cell (DC) subsets we needed to include CD141, FcER1, cDC1c and CD163 (8,9). CD141 typically shows trimodal distribution, with the CD141hi cells being cDC1s. We assigned CD141 to BUV615, which generates significant spreading error in the PE/Spark-YG593-area of the spectrum (both of which were assigned to mutually exclusive lineages).

For cDC2 identification we assigned FcER1 to BV711 and cDC1 to Pacific Blue, both of which delivered good to excellent resolution. For CD163 we tested PE-CF594, which as a fluorochrome was excluded from panel development early on, and APC-Cy7, which is a dim fluorochrome but delivered sufficient resolution to delineate CD163+ and – cells.

- PerCP was one of the three dimmest fluorochromes available in our selection. Thus we assigned it to CD11b, which can be used for subsetting NK cells, monocytes and DCs, and is typically expressed at relatively high levels.
- Zombie NIR and LIVE/DEAD™ Fixable Blue were both considered for the viability stain. Both fluorescent viability stains have a unique spectra and peak channel as compared to other fluorochromes commonly used for flow cytometry. AF350 and StarBright UV445 (two fluorochromes that we considered adding to the panel) do have substantial spectral similarity with LIVE/DEAD™ Fixable Blue (data not shown). Autofluorescence, additionally, has the same peak channel as LIVE/DEAD™ Fixable Blue and shared considerable spectral overlap. We therefore choose Zombie NIR for the viability stain due to the lower complexity number (CN) within the panel as compared to LIVE/DEAD™ Fixable Blue.

### Panel testing and validation

Initial panel testing involved a mixture of several smaller backbone panel runs both on the BD FACSDiscover™ S8 and the Sony ID7000™ (data not shown), a smaller 42-color panel (data not shown), as well as the testing of various fluorochrome-antibody conjugates using more recently released classes of fluorochromes, including NovaFluor dyes (formerly Phitonex, now available from Thermo Fisher), PE-Fire, APC-Fire and Spark conjugates (BioLegend), and RealBlue conjugates (BD Biosciences). Furthermore, we also tested various Qdot conjugates, which have long been available (10) but are now gaining attention again because of their relatively unique spectral profiles.

A long list of reagents was evaluated, but most were not included in the final panel. Below is a non-exhaustive list of reagents that were tested, but failed to perform for the stated reasons:

#### Reagents primarily excited by the UV laser

- Qdot 525 :: CD27 – The reagent was too dim to resolve CD27 intermediate staining. Qdot 525 was eliminated from the selection of fluorochromes.
- Qdot 585 :: CD8a – The reagent performed well. CD8a was switched to NovaFluor Blue 555 to accommodate spreading error between RB545, NovaFluor Blue 555, and NovaFluor Blue 585.
- Qdot 585 :: CD88 – The reagent required primary and secondary antibody staining due to limited commercial availability of directly conjugated Qdot 585 antibodies. This staining was unreliable. Qdot 585 was eliminated from the selection of fluorochromes.
- Qdot 655 :: CD134 – The reagent performed well. CD134 was eliminated from the selection of antigens due to low relevance to peripheral blood samples.
- Qdot 705 :: CD8a – The reagent performed well. CD8a was switched to NovaFluor Blue 555 as previously mentioned.
- Qdot 705 :: CX3CR1 – The custom conjugated reagent did not deliver the expected resolution. Qdot 705 was eliminated from the selection of fluorochromes.
- Qdot 705 :: HLA-DR – The reagent was too dim to resolve HLA-DR-high and HLA-DR-low populations. HLA-DR was switched to PE-Fire 810.
- Qdot 800 :: CD11c – The reagent was too dim to resolve DC population reliably. CD11c was switched to BV510.
- Qdot 800 :: CD19 – The reagent was too dim to reliably resolve the B cell population. Qdot 800 was eliminated from the selection of fluorochromes.
- Alexa Fluor 350 :: CD137 – The custom conjugate did not deliver the expected resolution. Alexa Fluor 350, LIVE/DEAD™ Fixable Blue Dead Cell Stain, and StarBright UltraViolet 445 were eliminated from the selection of fluorochromes to better model autofluorescence in similar spectral space (excitation 349 nm, emission 440 nm).
- LIVE/DEAD™ Fixable Blue Dead Cell Stain – The reagent performed well and had been used successfully in previous panels. The viability stain was switched to Zombie NIR Fixable Viability stain to better model autofluorescence as previously mentioned.
- StarBright UltraViolet 445 :: CD4 – The reagent performed well. StarBright UltraViolet 445 was eliminated from the selection of fluorochromes as previously mentioned.
- BUV496 :: CD16 – The reagent performed well and was successfully used in previous panels. Due to fluorochrome availability CD16 was switched to PE-AF700.
- BUV496 :: CD80 – The reagent did not deliver the expected resolution.
- BUV563 :: CD163 – The reagent did not deliver the expected resolution.\
- BUV661 :: CD137 – The reagent did not deliver the expected resolution.
- BUV661 :: CD38 – The reagent performed well. CD38 was substituted to APC/Fire 810 due to reagent availability and to prevent spreading error from BUV661 into other fluorochromes.
- BUV661 :: CD39 – The reagent performed well. CD39 was switched to R718 to prevent spreading error from BUV661 into other fluorochromes.
- BUV737 :: CX3CR1 – The reagent did not deliver the expected resolution.
- BUV805 :: NKG2C – The reagent did not deliver the expected resolution.

#### Reagents primarily excited by the violet laser

- BV421 :: CD25 – The reagent performed well. CD25 was switched to BB515 with similar separation so that CD161 could be used on BV421 to accommodate spreading error with Super Bright 436 :: IgM.
- BV421 :: GITR – The reagent performed well. GITR was removed from the selection of antigens to include other phenotyping markers.
- BV421 :: MR1-Tetramer (Ligand: 5-OP-RU) – The reagent performed well. Tetramers were omitted from further testing to incorporate the usage of Biotin-Streptavidin staining for fluorochromes with low commercial availability. TCR Va7.2 was used instead to delineate MAIT cells.
- eFluor 450 :: CD163 – The reagent performed well. eFluor 450 was eliminated from the the selection of fluorochromes in favor of Pacific Blue. CD163 was switched to APC-Cy7 in favor of Pacific Blue :: CD1c.
- BV480 :: CD5 – The reagent performed well. CD5 was eliminated from the selection of antigens to include other phenotyping markers.
- eFluor506 :: CD11c – eFluor506 was eliminated from the panel in favor of CD11c-BV510.
- BV510 :: CD20 – The reagent performed well. CD20 was switched to Spark YG 593 to accommodate commercial availability of antigens on Spark YG 593.
- Spark Violet 538 :: CD3 – The reagent provided insufficient resolution for CD3. Spark Violet 538 was eliminated from the selections of fluorochromes.
- BV570 :: CD45RA – The reagent performed well and used was successfully in previous panels. CD45RA was switched to Spark UV 387 to accommodate commercial availability of antigens on Spark UV 387.
- BV605 :: CD39 – The reagent worked well and was used successfully in previous panels. However, BV605 was assigned to CCR7, and thus CD39 was moved to R718 to with similar overall resolution.
- BV786 :: CD123 – The reagent worked well. CD123 was switched to PE-Cy5 to accommodate spread of PE-Cy5 into T cell phenotyping markers BB660 :: CD27, PE/Fire 640 :: CXCR3, APC :: TCR Va24-Ja18, AF647 :: TCR Va7.2.
- BV786 :: CD127 – The reagent performed well and was successfully used in previous panels. CD127 was switched to RB744 which had superior resolution of CD25 high, CD127low (Tregs).

#### Reagents primarily excited by the blue laser

- BB515 :: CD25 (Clone M-A251) – The reagent provided sub-optimal resolution for CD25. Clone BC96 BB515 :: CD25 had superior performance and was subsequentially used.
- BB515 :: CXCR5 – The reagent performed well. CXCR5 was eliminated from the selection of antigens to include other phenotyping markers.
- FITC :: CD8a. FITC was eliminated from the list of fluorochromes in favor of BB515 and to minimize the complexity number.
- Spark Blue 550 :: CD14 – The reagent performed well and was substituted with RB545 :: CD14 because of slightly better stain index.
- BB630 :: BTLA – The reagent performed well and was successfully used in previous panels. BB630 was eliminated in favor of RB613 and RB613 :: BTLA was used instead.
- BB630 :: CTLA4 – The reagent performed well and was successfully used in previous panels. CTLA4 was eliminated from the selection of antigens to prevent the need to perform intercellular staining.
- BB660 :: CD127 – The reagent performed well and was successfully used in previous panels. For reasons of commercial availability, CD127 was moved to RB744 with similar staining performance.
- BB660 :: PDL1 – The reagent performed well and was used successfully in previous panels. PDL1 was eliminated from the selection of antigens due to limited relevance to peripheral blood samples.
- PerCP :: CD5 – The reagent performed well. CD5 was eliminated from the selection of antigens to include other phenotyping markers.
- PerCP :: CD8a – The reagent performed well. CD8a was switched to NovaFluor Blue 555 as previously mentioned.
- BB700 :: CD161 – The reagent performed well and was used successfully in previous panels. CD161 was moved to BV421 and BB700 :: IgG was used instead to accommodate spread of BB700 into APC :: TCR Va24-Ja18 and AF647 :: TCR Va7.2.
- BB700 :: CD32 – The reagent performed well and was used successfully in previous panels. CD32 was eliminated from the selection of antigens to include other phenotyping markers.
- PerCP-eFluor710 :: TCRgd – The reagent worked well and was successfully use in previous panels (OMIP-069). PerCP-eFluor710 was eliminated from the selection of fluorochromes in favor of BB700 and RB705.
- BB755 :: CD28 – The reagent worked well. BB755 was eliminated in favor of RB744. CD28 switched to BV480 to accommodate spreading error between BV480 and AmCyan :: CD19 in addition to BV480 and Pacific Blue :: CD1c.
- BB755 :: Granzyme B – The reagent performed well. Granzyme B was eliminated from the selection of antigens to prevent the need to perform intercellular staining.
- BB790 :: TIGIT – The reagent performed well and was successfully used in previously published panels. BB790 was eliminated in favor of RB780. In comparison testing, RB780 :: TIGIT (clone: TgMab-2) was shown to outperform RB780 :: TIGIT (clone: 741182).

#### Reagents primarily excited by the yellow-green laser

- PE :: IL18R1 – The reagent provided insufficient resolution of IL18R1. IL18R1 was eliminated from the selection of antigens due to low relevance in peripheral blood samples.
- Spark YG 593 :: CD19 – The reagent performed well. CD20 was switched to Spark YG 593 and CD19 was moved to AmCyan to accommodate commercial availability of antigens on AmCyan.
- NovaFluor Yellow 590 :: CD19 – The reagent performed well. NovaFluor Yellow 590 was eliminated from the selection of fluorochromes in favor of Spark YG 593 which had less background staining.
- PE-CF594 :: CD163 – The reagent performed well and has been used successfully in previous panels (OMIP-044). PE-CF594 was eliminated from the selection of fluorochromes in favor of PE-AF610. CD163 switched to APC-Cy7 with sufficient resolution.
- PE-CF594 :: TCRgd – The reagent performed well and has been used successfully in previous panels. The fluorochrome PE-CF594 eliminated in favor of PE-AF610. TCRgd switched to RB705 to accommodate spreading error with BB700 :: IgG.
- PE/Fire 640 :: CD45RO – The reagent performed well, but CD45RO was switched to BV570, performing equally well.
- PE-Cy5 :: CD80 – The reagent performed well and was used successfully in previous panels (OMIP-044). CD80 was eliminated from the selection of antigens to include other phenotyping markers.
- PE-Cy5.5 :: CD19 – The reagent performed well and has been used successfully in previous panels (OMIP-044). CD19 was switched to AmCyan to accommodate commercial availability of antigens on AmCyan.

#### Reagents primarily excited by the red laser

- Alexa Fluor 647 :: CD1c – The reagent worked well and was used successfully in previous panels (OMIP-044). Due to unmixing spreading error greatly dimensioning the resolution of Alexa Fluor 647, a more highly expressed, bi-modal antigen density target was desired (in this case TCR Va7.2). CD1c was switched to Pacific Blue.
- Spark NIR 685 :: CD123 – The reagent performed well. Due to unmixing spreading error CD123low basophils were not easily resolved. CD123 was switched to PE-Cy5.
- Spark NIR 685 :: CD19 – The reagent performed well and was used successfully in previous panels (OMIP-069). CD19 was switched to AmCyan to accommodate commercial availability of antigens on AmCyan.
- Spark NIR 685 :: CD27 – The reagent performed well, but CD27 was kept at a previously used conjugate because of limited availability (BB660 :: CD27).
- Spark NIR 685 :: CXCR5 (CD185) – Spark NIR 685 is one of the fluorochromes most affected by unmixing spreading error, and as a result separation was insufficient to resolve CXCR5+ cells. Marker was dropped in favor of a more highly expressed T cell target (KLRG1).
- Alexa Fluor 660 :: CD56 – Alexa Fluor 660 suffered from significant unmixing spreading error and delivered insufficient resolution to separate CD56bright and CD56dim NK cells.
- Alexa Fluor 660 :: CD19 – As mentioned, Alexa Fluor 660 suffered from significant unmixing spreading error. Alexa Fluor 660 was eliminated from the selection of fluorochromes in favor of Spark NIR 685.
- NovaFluor Red 660 :: CD4 – The regent worked well. NovaFluor Red 660 was eliminated from the selection of fluorochromes in favor of APC.
- NovaFluor Red 700 :: CD8a – The regent worked well. NovaFluor Red 700 was eliminated from the selection of fluorochromes in favor of Spark NIR 685.
- APC-R700 :: CCR7 (CD197) – The reagent performed well. Due to fluorochrome availability, CCR7 (CD197) was moved to BV605. APC-R700 was subsequently eliminated from the selection of fluorochromes in favor of R718.
- APC-H7 :: CD45RA – The reagent performed well. CD45RA was switched to Spark UV 387 due to commercial availability of antigens on Spark UV 387. APC-Cy7 was used instead of APC-H7 in the final fluorochrome selection.
- Alexa Fluor 790 :: CD206 – The custom conjugate did not provide the expected resolution. Alexa Fluor 790 was removed from the selection of fluorochromes in favor of APC/Fire 810.
- APC/Fire 810 :: CD45 – The reagent performed well. CD45 was moved to BUV805 to accommodate CD38 on APC/Fire 810.

For the final and optimized panel, we ran 10 individual fluorescence-minus-one (FMO) controls to formally exclude potential spreading-error problems for markers that were considered of high importance. Some of these FMO controls are shown in the main Figure 1, while the remaining FMO controls are depicted in **Online Figure 9A**. Furthermore, for all markers in the panel we evaluated staining specificity by cross-checking expression patterns across the main immune cell lineages (T cells, B cells, NK cells, DCs, monocytes). A selection of these validation histograms for phenotyping markers that did not have a dedicated FMO control are shown in **Online Figure 9B**. Also, we designed a “trimmed-down” 45-color panel that we used primarily to evaluate how the addition of the 5 fluorochromes affects unmixing spreading error and overall panel performance relative to the full 50-color panel (data not shown).

We also leveraged this 45-color panel as a “fluorescence-minus-5” control, again showing that all the 5 fluorochromes added to the 50-color panel deliver good separation, without any issues from SE (**Online Figure 9C)**. Notable, unmixing the 45-color panel with a 45-color unmixing matrix would have resulted in reduced unmixing spreading error (i.e. reduced spread and tighter negative populations) for several fluorochromes (data not shown).

**Online Figure 9:**
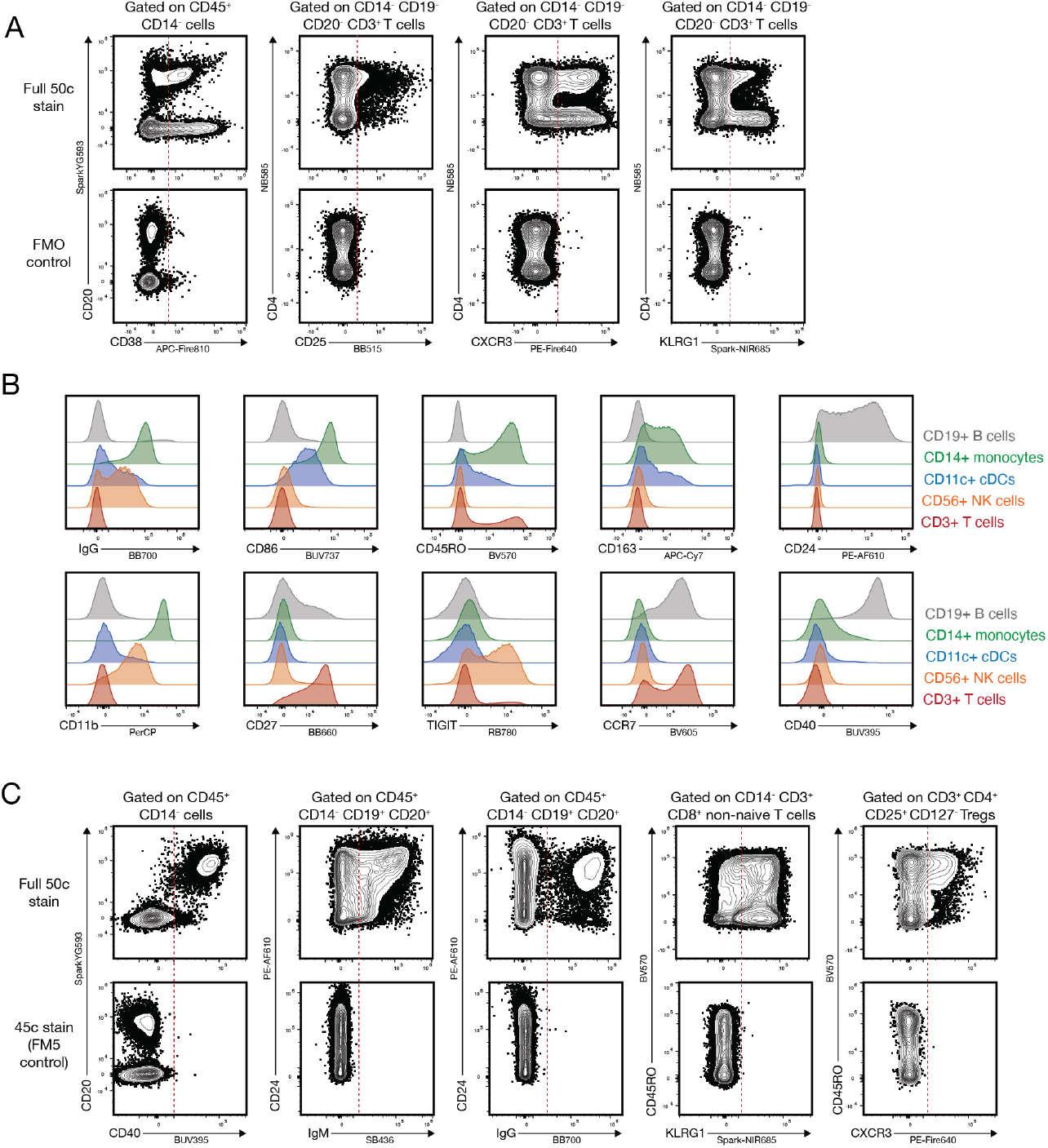
Additional FMO controls and gating controls. PBMCs were obtained from commercial vendors, stained with the full 50-color (50c) or the trimmed-down 45-color (45c) panel and acquired on a BD FACSDiscover™ S8. (A) Additional fluorescence-minus-one (FMO) controls for the indicated markers: CD38, CD25, CXCR3 and KLRG1. Pre-gating is indicated above the plots. Dotted red lines indicate positivity cut-offs. (B) Histogram overlays for the indicated markers on B cells (grey), monocytes (green), cDCs (blue), NK cells (orange) and pan T cells (red). (C) Plots from the full 50c stain (top panel) and trimmed-down 45c panel (lower panel) lacking the indicated reagents. For the plots in panel (C) OLS unmixing data is shown. Dotted red lines indicate positivity cut-offs. Pregating is indicated above the plots.

Moreover, in the full 50-color panel we visually evaluated the “most problematic” fluorochrome pairs based on high similarity index and known fluorochrome properties (**Online Figure 10**). As expected, these pairs showed some degree of tilted appearance of the negative population, as has been described both for conventional compensation-based and spectral cytometry before (11). However, for all these problematic pairs separation of the stained markers was good enough to demarcate the positive populations, even without any major pregating.

**Online Figure 10:**
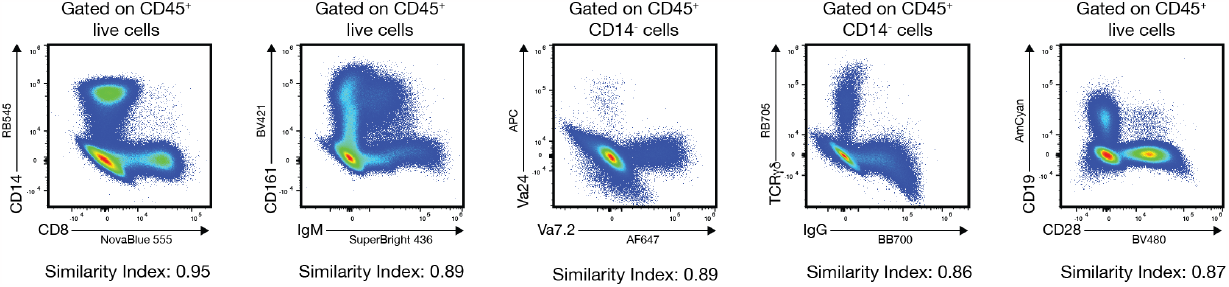
Fluorochrome pairs with high similarity index. PBMCs were obtained from commercial vendors, stained with the full 50-color panel and acquired on a BD FACSDiscover™ S8. Plots depict either all CD45+ live events or CD45+ non monocyte events, as indicated above plots. These fluorochrome pairs showed the highest similarity indices in our fluorochrome selection (see Online Figure 3), but still resolve well with relatively low spreading error (SE).

Finally, we also evaluated whether our panel performs well consistently across multiple donors for all markers of a populations of interest. We stained different donors and thoroughly assessed staining patterns and resolutions across the entire panel. As expected, for lineage markers, positive and negative populations remained clearly separated, with the expected minimal stain-to-stain-variability (data not shown). More importantly, the resolution for both the T cell phenotyping markers and APC phenotyping markers remained very good across multiple donors. **Online Figure 11** shows histogram overlays for these markers across three different donors.

Of note, all the raw data acquired on the BD FACSDiscover™ S8 (main Figure 1 and the online figures depicting gating controls) has been deposited on Flowrepository with the identifier <u>FR-FCM-Z73V</u> and is publicly available for download.

#### Cross-platform panel

As mentioned in the main text and introduction, one of our goals was to develop a panel that is not only optimized to one given instrument (in this case the BD FACSDiscover™ S8), but also cross-platform compatible across at least one additional spectral cytometry platform, in this use case the Sony ID7000™.

The optical setup of the Sony ID7000™ had the same excitation laser wavelengths as the BD FACSDiscover™ S8 and in addition a near UV laser (320nm) and Near-Infrared laser (808nm). Initial tests showed that the usage of the NIR laser was not beneficial because of the automatic addition of a Notch filter for the same wavelength range across the other detector arrays (data not shown). Thus, we decided to perform all our measurements on the Sony ID7000™ using a 6-laser setup with the 808nm laser switched off.

**Online Figure 11:**
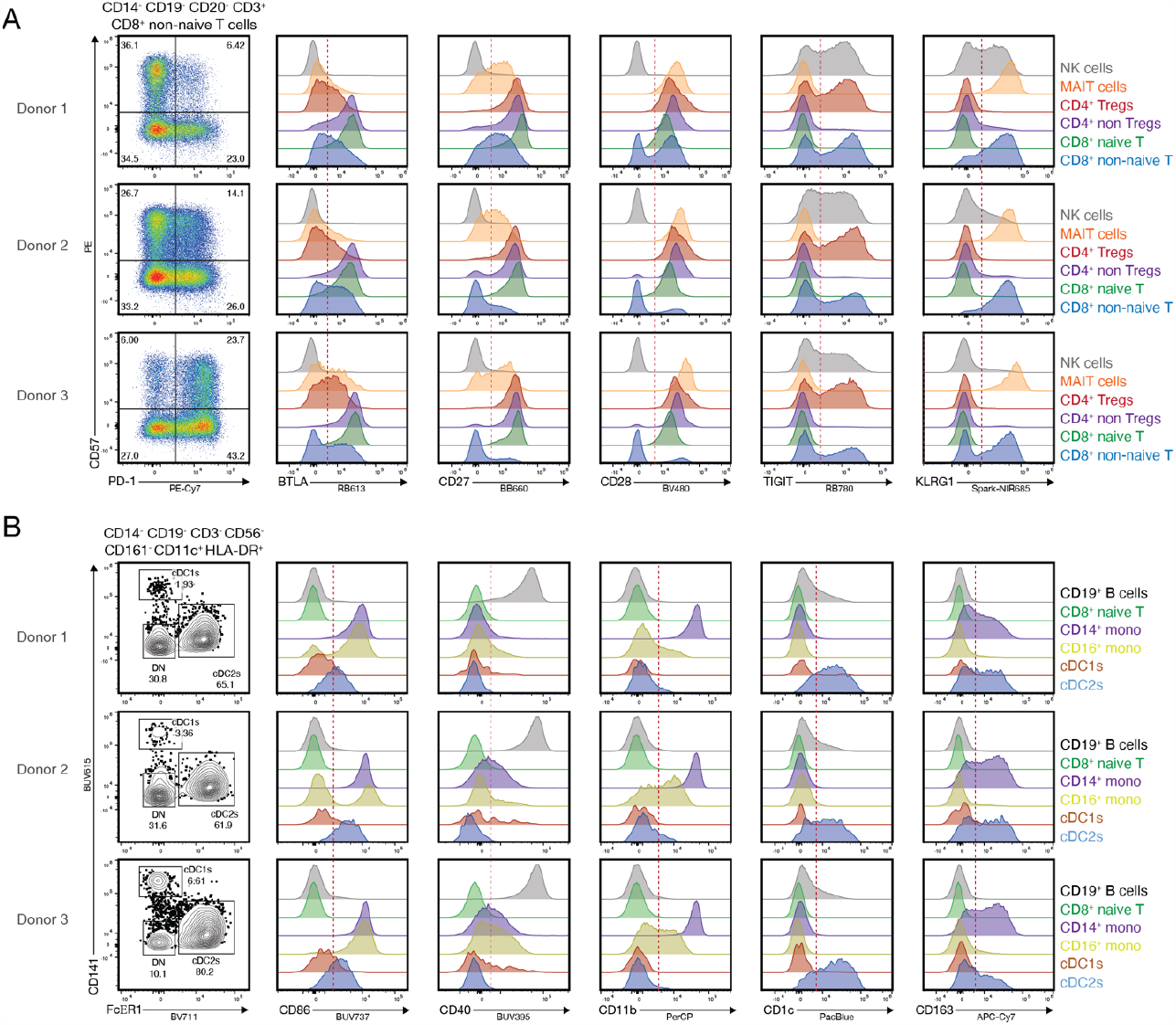
Consistency in resolution across multiple donors. PBMCs were obtained from commercial vendors, stained with the full 50-color (50c) panel and acquired on a BD FACSDiscover™ S8. Three different donors are shown. (A) 2D plots for CD57 and PD-1 on the CD8+ non-naïve T cell population (left side, gated as in main Figure 1) and histogram plots for the five indicated markers across the listed T cell populations. (B) 2D plots for CD141 and FcER1 on the CD11c+ HLA-DR+ cDC population (left side, gated as in main Figure 1) and histogram plots for the five indicated markers across the listed B cell, monocyte and DC populations. Red dotted lines show positivity cut-offs.

With respect to detection optics, the Sony ID7000™ collects at least the same range of emission wavelengths, but using a different detector layout based on a 32-channel PMT with more narrow detection windows, and thus an overall higher number of detector bands (12). However, throughout our panel development we observed that certain spectrally similar fluorochromes after unmixing showed such high levels of spectral unmixing error (i.e. noise) that there would be very limited or no possibility for meaningful resolution of biological targets. These pairs were: BV421 and Super Bright 436, SparkUV387 and BUV395, BB700 and RB705, PE-Fire 640 and other dyes in the same emission area, Spark NIR 685 and the neighboring dyes in the same emission area.

**Online Figure 12:**
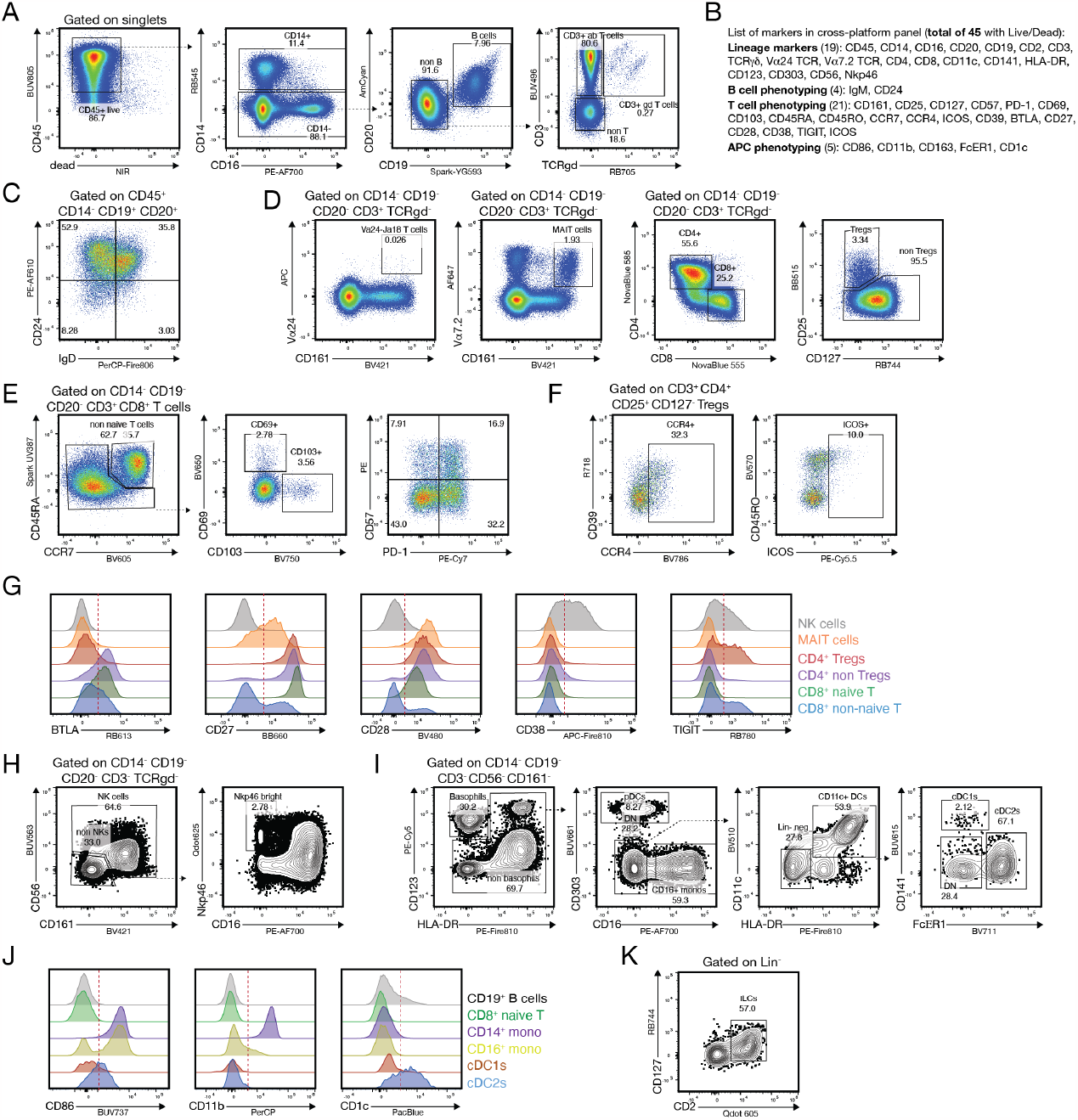
Cross-platform compatible 45-color panel acquired on a Sony ID7000™. PBMCs were obtained from commercial vendors, stained as described in the online section of the manuscript and acquired on Sony ID7000™. The detector gains on the Sony ID7000™ apply to an entire detector array devoted to an individual laser. Detector gains were determined via a voltration-like method and set where a select group of fluorochromes on CD4 for a given laser detector array all had optimized stain indexes, or to the highest setting while keeping the brightest CD4 fluorochrome on scale. Pre-gating of plots is annotated in the figure or indicated by dotted black arrows. For some plots different donors are shown. Abbreviations: BUV: Brilliant Ultraviolet; BV: Brilliant Violet; BB: Brilliant Blue; RB: RealBlue; AF: AlexaFluor; PE: Phycoerythrin; APC: Allophycocyanin; Qdot: Quantum Dot; NIR: Near-Infrared; The overall gating strategy is copied from the one used for the full 50c panel shown in Figure 1. (A) Gating strategy for CD45+ live cells, monocytes, B cells and γδ and αβ T cells. (B) Overview of the 45 targets analyzed with this cross-platform compatible panel. (C) Representative plots for the IgD and CD24 expression on B cells. (D) Gating strategy to delineate invariant NKT cells, MAIT cells, CD4+ and CD8+ T cells, as well as CD4+ regulatory T cells (Tregs).(E) Representative plots for CD69, CD103, CD57 and PD-1 expression on non-naïve CD8+ cytotoxic T cells. (F) Expression patterns for CD39, CCR4, CD45RO and ICOS on the CD4+ Treg population. (G) Histogram overlays for the expression pattern of BTLA, CD27, CD28, CD38 and TIGIT on NK cells (grey), MAIT cells (orange), CD4+ Tregs (red), CD4+ non Tregs (purple), CD8+ naïve T cells (green) and CD8+ non-naïve T cells (blue). Dotted red lines indicate positivity cut-offs. (H) Gating strategy for NK cells and NK cell subsets based on CD16 and Nkp46. (I) Gating strategy for Basophils (CD123+ FcER1+), plasmacytoid DCs (CD303+ HLA-DR+), pan conventional DCs (CD11c+ HLA-DR+), and the cDC1 (CD141+) and cDC2 (FcER1+) subsets. (J) Histogram overlays for the expression pattern of CD86, CD11b, and CD1c on B cells (grey), CD8+ naïve T cells (green, negative control), CD14+ monocytes (purple), CD16+ monocytes (yellow), CD141+ cDC1s (red) and FcER1+ cDC2s (blue). Dotted red lines indicate positivity cut-offs. (K) Gating strategy for Lin-CD2+ CD127+ innate lymphoid cells (ILCs).

Thus, we decided to omit the phenotyping markers IgM-Super Bright 436, IgG-BB700, CD40-BUV395 (to keep the more relevant CD45RA), CXCR3-PEFire 640 and KLRG1-Spark NIR 685 for a trimmed-down panel version that we ran on both the BD FACSDiscover™ S8 and Sony ID7000™. For the Sony ID7000™, data was unmixed using the weighted least-square (WLSM) method, which is the default unmixing method in the Sony acquisition software.

**Online Figure 12** shows the same gating strategy as in main Figure 1 (minus the removed five markers mentioned above). Of note, we do not intend to compare instrument performance in detail, but rather would like to highlight that a complex panel can be designed in such a way that it can be used on two independent spectral cytometers to facilitate cross-platform studies. To the best of our knowledge this is the first report of a 45-color phenotyping panel that has been tested to be cross-platform compatible. Future work is needed to test additional platforms from other vendors and to systematically characterize the resolution of individual fluorochromes in the context of very complex unmixing matrices across these platforms.

#### Spectral unmixing and reference controls

With more high-dimensional spectral cytometry data and best practices being publicly available, it is becoming evident that the quality of single-stained controls is the most critical aspect of the success for such a panel. Post-acquisition correction of unmixing matrices to correct apparent unmixing errors is in the best case very cumbersome, and in the worst case a source for artifacts.

Thus, with multiple panel runs, we generated 50 single-stained controls with cells (PBMCs) and antibody capture beads (BD Compbeads, see staining protocol section below). We carefully evaluated differences in emission spectra for all fluorochromes and observed results similar to previous reports where some fluorochromes show slight changes in emission when bound to polystyrene beads compared to cells (**Online Figure 13**). This highlights that for the success of a complex high-dimensional panel the choice of single-stained control is absolutely critical to achieve correct unmixing. For our final unmixing matrix, we exclusively used the cell-based controls as input. Future studies are needed to build a comprehensive database listing which fluorochrome spectra vary between different types of capture beads (polystyrene or cell-mimics) and cells.

To identify potential unmixing artifacts, we carefully checked NxN plots of all fluorochrome combinations in our final panel (data not shown for space reasons). Of note, we found only a handful of inaccuracies that did not affect the biological expression patterns of our data. Thus, we decided to not perform any adjustments of the unmixing matrix on this data set.

**Online Figure 13:**
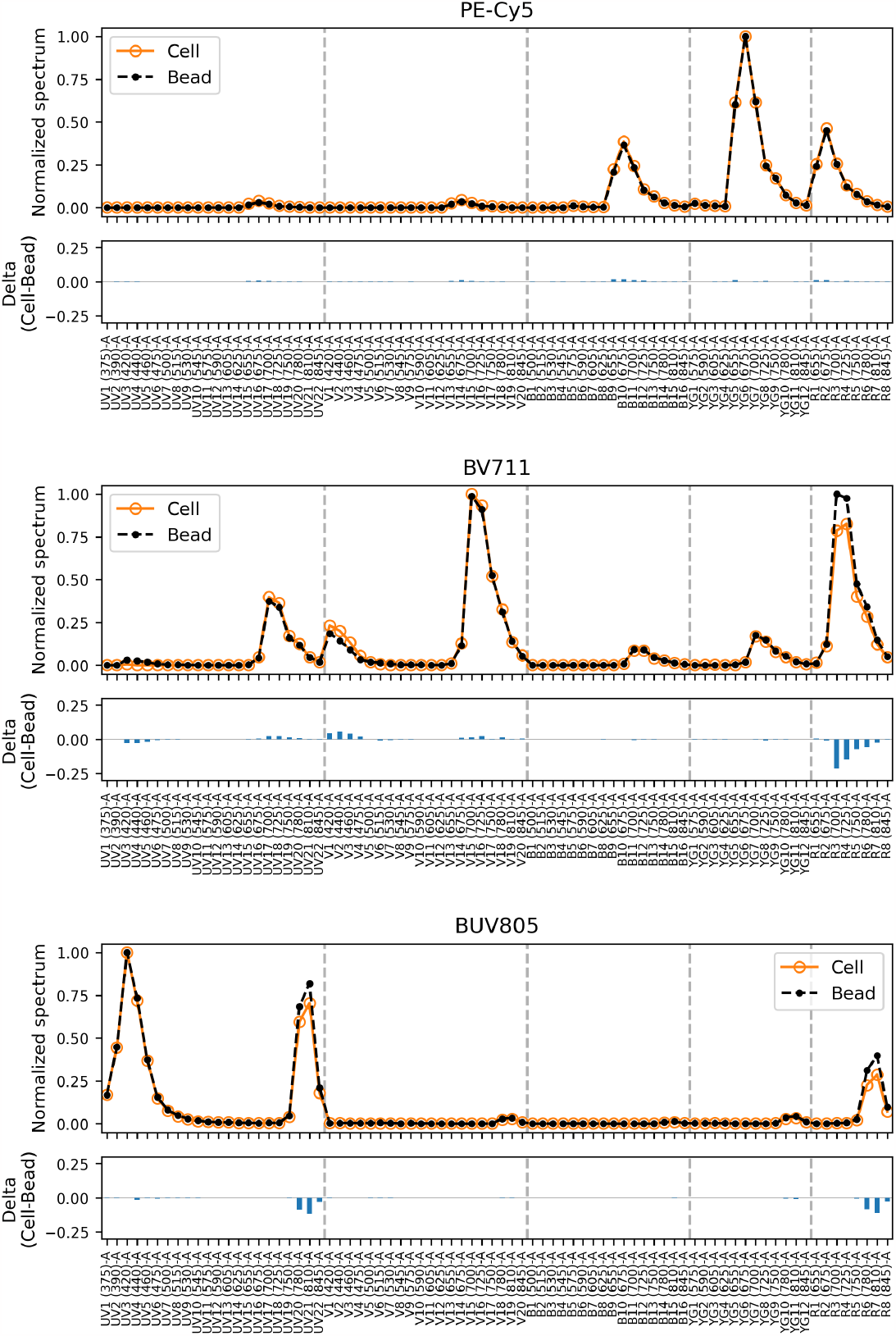
Example comparisons of single-stain spectra for beads and cells. Reagent-matched single-stain controls were prepared using both PBMCs and BD Comp Beads and acquired on a BD FACSDiscover™ S8 at standard settings. Normalized spectra match closely between beads and cells for most fluorochromes (PE-Cy5 shown as an example, top panel), but can differ significantly for other fluorochromes (BV711 and BUV805 shown as examples, center and bottom panels, see Delta Cell-Bead).

#### Cell sorting

Given the functionality of the BD FACSDiscover™ S8 as a cell sorter (1), we evaluated whether the 50-color panel was also suitable for cell sorting.

**Online Figure 14:**
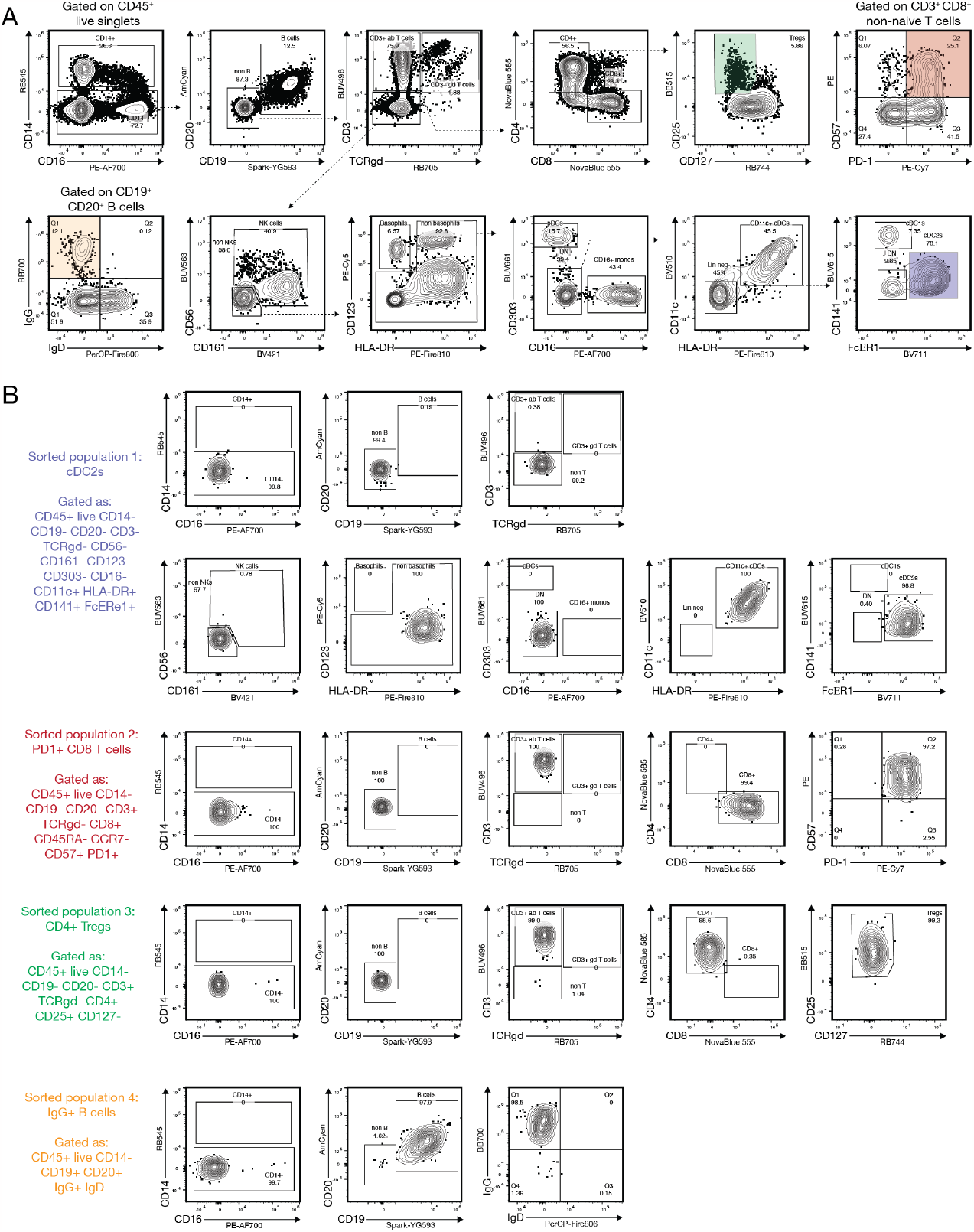
Example sorting results using the full 50-color panel. PBMCs were obtained from commercial vendors, stained with the 50-color panel and sorted for four different populations on a BD FACSDiscover™ S8. Instrument setup for cell sorting followed best practices and the instructions from the manufacturer. The instrument was operated at a sheath pressure of 35psi with an 85um nozzle. (A) Gating strategy to delineate the main immune cell subsets and all populations that were planned for sorting. Sorted populations are marked with colored boxes. (B) Re-analysis of the indicated populations post sorting. Color-code referring to the gating tree.

We chose 4 different target populations to cover relatively fine-grained and rare phenotypes as well as more abundant cell types: CD57^+^ PD1^+^ non-naïve CD8^+^ T cells, CD4^+^ Tregs, FcER1^+^ cDC2s and IgG^+^ B cells. **Online Figure 14** shows the original sample sorted and gating strategy and the post-sort purity for each of the gated populations (**Online Figure 14B**). This highlights that our panel can be used to generate 50-color phenotyping data while successfully sorting on distinct cell populations as needed. The ability to generate parallel high-dimensional phenotyping data while sorting can be particularly useful for limited human tissue samples.

#### Computational analysis of data

In many cases, high-dimensional cytometry panels are utilized for exploratory data analysis (13,14). As anticipated, our 50-color panel is able to resolve heterogeneity in the immune compartment in a detailed manner, particularly in the T cell and APC compartment. Many computational tools are available to perform this cellular heterogeneity, e.g. dimensionality reduction and clustering (PhenoGraph, FlowSOM, and others), or full annotation using-shape-constrained trees (FAUST). In **Online Figure 15** we show a representative dimensionality reduction plot using uniform manifold approximation projection (UMAP), overlaid with 30 FlowSOM meta clusters for the entire CD45+ live population. As can be seen in the heatmap with scaled marker expression (**Online Figure 15B**), even low-abundance populations such as γδ T cells or CD141+ cross-presenting cDC1s are identified as unique clusters. Reclustering the data after subsetting on either T cells or APCs results in further defined heterogeneity (data not shown).

Of note, **Online Figure 15C** shows histogram overlays for some lineage markers (CD20, CD3, CD14), as well as representative functional phenotyping markers for the T cell compartment (CD25, TIGIT, CD38) and the APC compartment (CD86 and CD163). These histogram overlays demonstrate that across all clusters there is consistent tight distributions of the negative population as well as distinct separation of the positive signals. This representation supports the notion that our 50-color panel delivers good resolution across all the cells in the sample.

**Online Figure 15:**
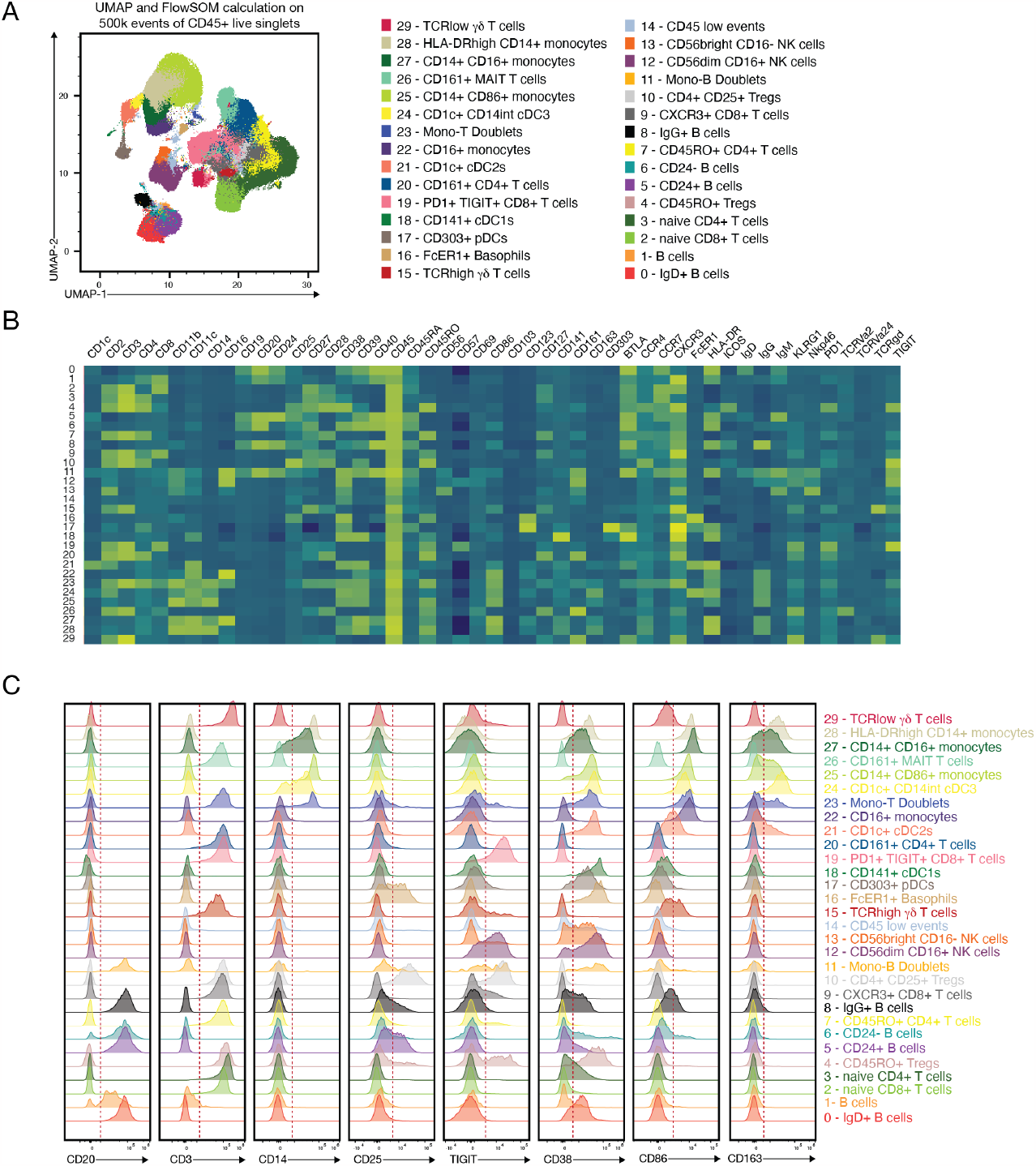
Exploratory analysis using dimensionality reduction and clustering. Data from the full 50-color panel acquired on the BD FACSDiscover™ S8 was gated on CD45+ live events, subsampled to 500.000 events and subjected to UMAP and FlowSOM calculation, with 30 metaclusters. (A) FlowSOM metaclusters overlaid onto a UMAP plot with manual annotation of populations. (B) Heatmap overview of marker expression (scaled) for all FlowSOM clusters. (C) Histogram representation of the indicated markers across all FlowSOM clusters. Dotted red line indicates manual positivity cut-offs. Note the consistent and tight distribution of negative peaks across almost all clusters.

### Titrations

All reagents used in panel development have been obtained from commercial vendors (BD Biosciences, BioLegend, Thermo Fisher), as listed in Online Table 2 below. For all antibodies, we performed serial dilution series from 1:5 down to 1:640, or with additional dilutions as required (final staining volume of 50ul for 2-5×10^6^ cells). We performed co-staining either with CD3 or CD14, with pre-gating as required to evaluate the biological expression pattern of the titrated reagent. For all titrations, we calculated the stain indices and selected the best concentration either based on ideal stain index, or on the best possible resolution while limiting the signal intensity to minimize spreading error (e.g. for CD57).

**Online Figure 16** shows the concatenated titration plots for all reagents primarily excited by the UV, and violet laser. **Online Figure 17** shows the concatenated titration plots for all reagents primarily excited by the blue, yellow-green and red laser.

### Staining protocol

**Online Figure 16:**
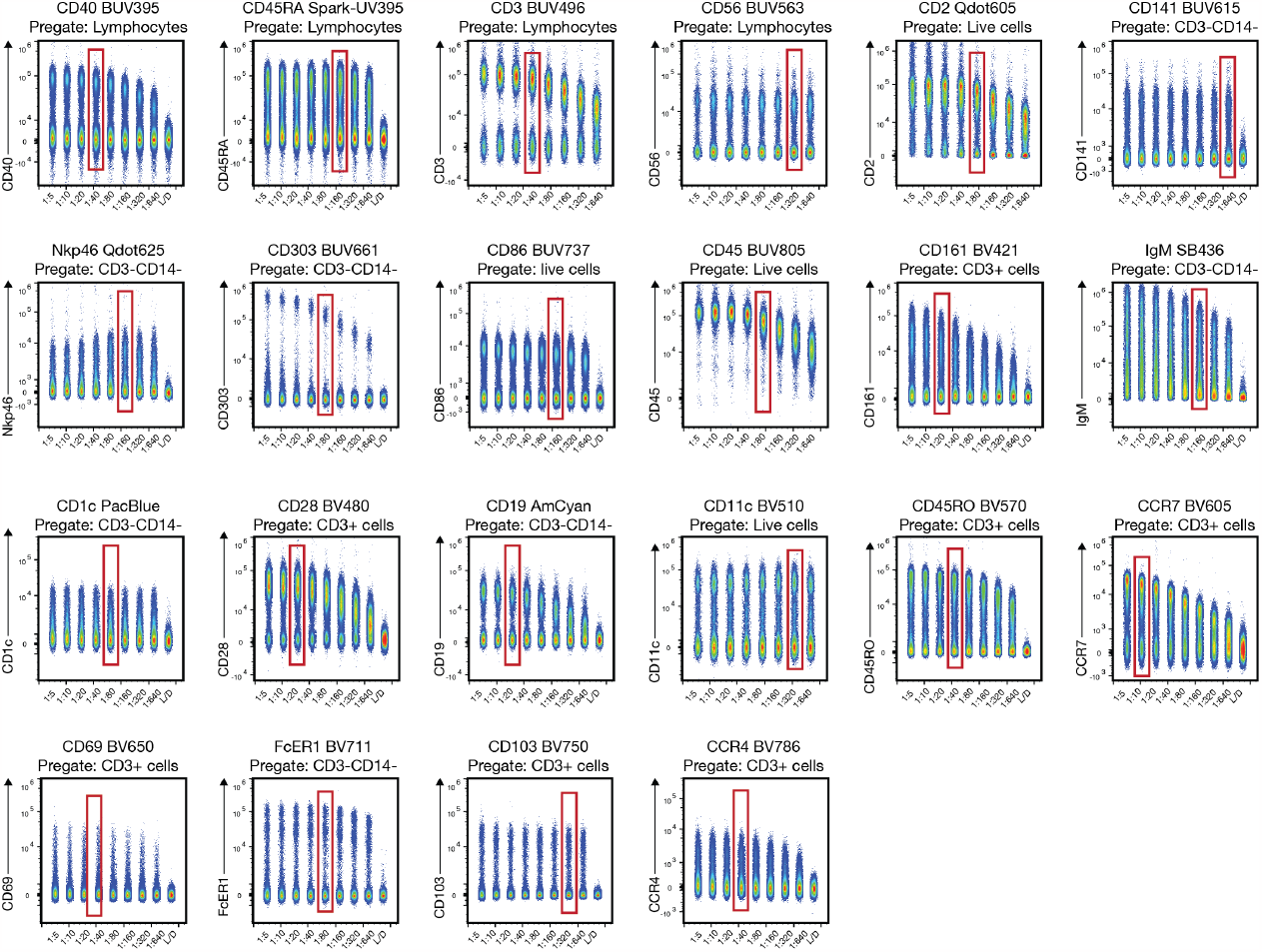
Titrations for fluorochrome-antibody conjugates primarily excited by the UV and violet laser. Titrations were performed on cryopreserved PBMCs using serial dilutions, starting at a highest concentration of 1:5 down to 1:640 (unless indicated otherwise) and acquired on the Sony ID7000™. Some antibodies were titrated in pairs of two or three (when no or minimal spectral overlap was present). All samples were pre-stained with CD3 and CD14, and the population used for evaluating the titration is indicated in the respective plot. Red squares depict the chosen concentration. Individual FCS files were concatenated for representation. L/D: only stained with L/D reagent (only included for some samples).

**Online Figure 17:**
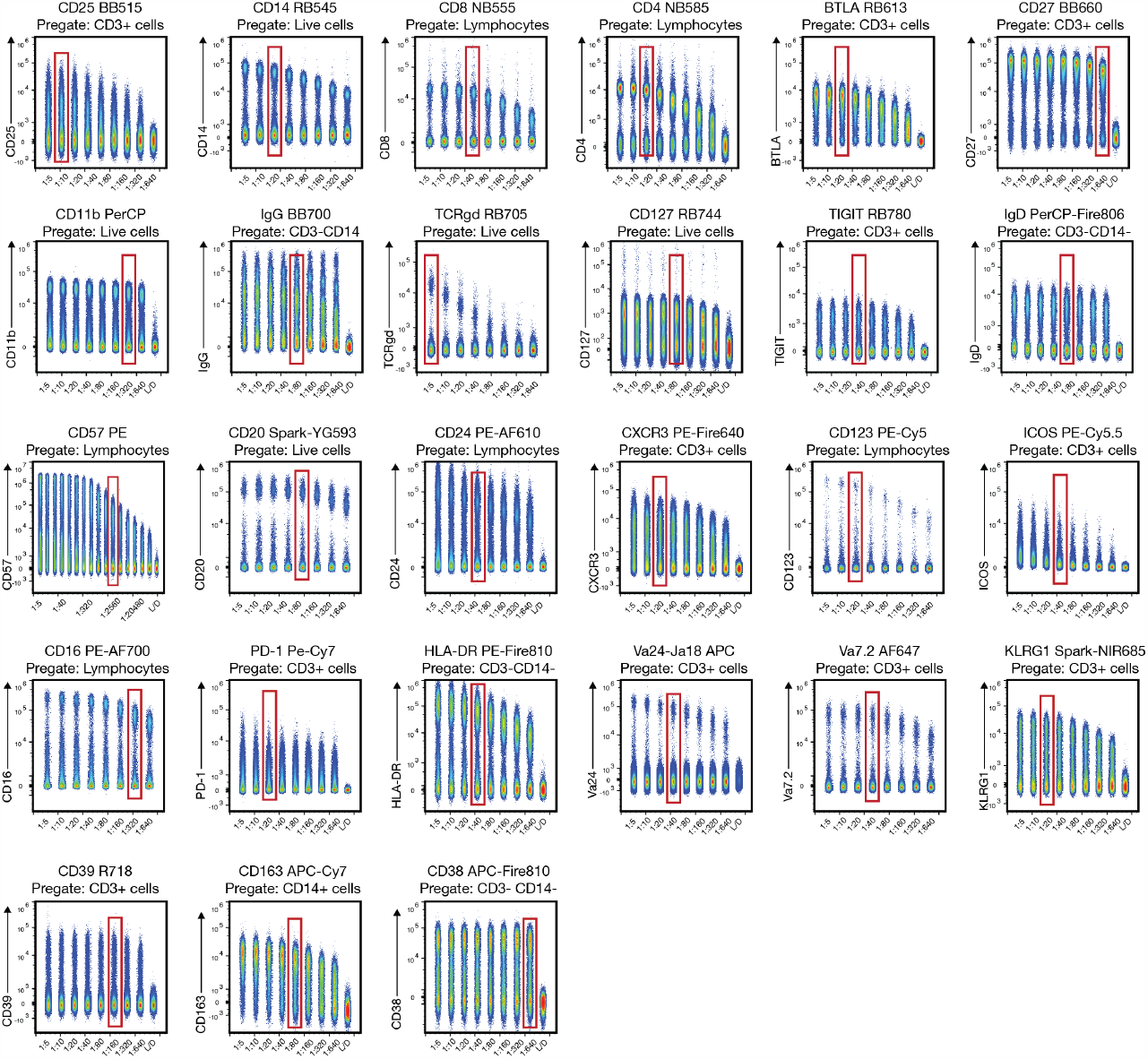
Titrations for fluorochrome-antibody conjugates primarily excited by the blue, yellow-green and red laser. Titrations were performed on cryopreserved PBMCs using serial dilutions, starting at a highest concentration of 1:5 down to 1:640 (unless indicated otherwise) and acquired on the Sony ID7000™. Some antibodies were titrated in pairs of two or three (when no or minimal spectral overlap was present). All samples were pre-stained with CD3 and CD14, and the population used for evaluating the titration is indicated in the respective plot. Red squares depict the chosen concentration. Individual FCS files were concatenated for representation. L/D: only stained with L/D reagent (only included for some samples).

**Online Figure 18:**
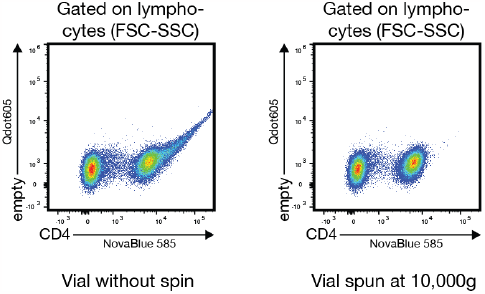
Removal of antibody aggregates by centrifugation. Cryopreserved PBMCs were stained with anti-CD4-NovaBlue 585 either with or without centrifugation of the antibody vial at 10.000g. Samples were acquired on a Sony ID7000™.

#### Commercial reagents

- RPMI Medium 1640 [1x] (Gibco #11875-093)
- FBS [heat-inactivated] (Peak Serum #PS-FB2)
- GlutaMAX-I [100x] (Gibco #35050-061)
- Penicillin-Streptomycin [10,000 U/mL] (Gibco #15140-163)
- DPBS (Gibco #14190-144)
- Zombie NIR Fixable Viability Kit (Biolegend #423106)
- Human TruStain FcX (Biolegend #422302)
- BD Horizon Brilliant Stain Buffer Plus (BD #566385)
- CellBlox Blocking Buffer (Invitrogen #H001T03B04)
- True-Stain Monocyte Blocker (Biolegend #426103)
- BD Cytofix/Cytoperm Fixation and Permeabilization Solution (BD #554722)
- BD CompBeads Anti-Mouse Ig, κ/Negative Control Compensation Particles Set (BD #552843)
- BD CompBead Plus Anti-Mouse Ig, κ/Negative Control (BSA) Compensation Plus (7.5 µm) Particles Set (BD #560497)
- BD™ CompBeads Anti-Rat and Anti-Hamster Ig κ /Negative Control Compensation Particles Set (BD #552845)
- ArC Amine Reactive Compensation Bead Kit (for use with LIVE/DEAD Fixable dead cell stain kits) (Invitrogen #A10346)
- Cryopreserved PBMCs were obtained from whole blood or leukopak purchased from Bloodworks Northwest (https://www.bloodworksnw.org/, Washington USA). Different donors were used throughout the panel development process.

#### Prepared buffers and procedure notes

- Cell culture media (RP10): add 50 ml of FBS, 5ml of GlutaMAX-I and 5 ml of Penicillin-Streptomycin into 500 ml of RPMI Medium 1640
- FACS buffer: add 10 ml of FBS to a 500 ml bottle DPBS.
- Fc-Block/Viability dye solution: add 25 ul of Human TruStain FcX™ and 1 ul of reconstituted Zombie NIR Fixable Viability reagent to 500 ul of DPBS immediately prior to use.

Antibody staining mix 1, Antibody staining mix 2, and FMO staining mixes: All antibody staining mixes are made on the day of their usage with a final volume of 50 ul per sample. The full panel is divided into 2 staining mixes to maintain the optimal concentration of fluorochromes as derived from titrations, to facilitate staining that require 2 steps, and to prevent documented interactions of steric hinderance.

Table # illustrates the correct final dilution of each antibody and to which antibody mix it has been assigned. To facilitate staining that requires 2 steps, Biotin :: Nkp46 (CD335) was assigned to antibody staining mix 1 and Qdot 625 :: Streptavidin was assigned to antibody staining mix 2. To prevent loss of staining to steric hinderance, RB705 :: TCRgd, APC :: TCR Va24-Ja18, and Alexa Fluor 647 :: TCR Va7.2 were assigned to antibody staining mix 1 and BUV496 :: CD3 was assigned to antibody staining mix 2. Targets that required FMOs were consolidated into antibody staining mix 2 to reduce the number of unique antibodies staining mixes and pipetting. All mixes contain Brilliant Stain Buffer Plus and True-Stain Monocyte Blocker at the manufacture’s recommended concentration to prevent aggregation of polymer dyes and non-specific staining of dyes containing cyaninine or cyanine-like molecules (15). Antibody staining mix 2 and FMO staining mixes contain CellBlox Blocking Buffer at the manufacture’s recommended concentration to prevent non-specific staining of NovaFluor dyes.

Reagents used during and after staining with Qdot reagents (FACS buffer, Brilliant Staining Buffer Plus, BD Cytofix/Cytoperm™ Fixation and Permeabilization Solution) were tested for heavy metal contamination to prevent Qdot quenching as has been prior reported. In short, single stain controls were made for Qdot 605 :: CD2 where 1 mM EDTA final concentration was added to each buffer during and after the staining process or not. The MFI of Qdot 605 was compared between the two conditions to determine whether quenching was taking place due to contamination (16). Only non-contaminated reagents were used without the addition of EDTA going forward (data not shown).

Antibody mixes were centrifuged for 10,000 x g for 5 minutes immediately prior to use to remove antibody aggregates. We have observed a decrease in aggregation of Qdot 605 :: CD2, BUV615 :: CD141, Qdot 625 :: NKp46 (CD335), Pacific Blue :: CD1c, NovaFluor Blue 555 :: CD8a, NovaFluor Blue 585 CD4, and RB744 :: CD127. An example for such aggregates before and after centrifugation is shown in **Online Figure 18**.

#### Step-by-step staining procedure

**Online Table 2:** Reagents used in the final panel. List of all 50 antibodies and Streptavidin used in the final panel. For each fluorochrome, we list the peak detector on the BD FACSDiscover™ S8 (third column). Titers refer to final dilution factors, and the final staining volume per sample was 50ul.

|  | Laser: | Fluorochrome: | BD FACSDiscover™ | Target: | Clone: | Vendor: | Catalog: | Titer: | Stain: |
| --- | --- | --- | --- | --- | --- | --- | --- | --- | --- |
| 0 |  | Biotin | NA | NKp46 | 9E2 | Biolegend | 331906 | 1:160 | 1 |
| 1 | 349 nm | Spark UV 387 | UV01 [375] (0.98),<br>UV03 [420] (1.00) | CD45RA | HI100 | Biolegend | 304179 | 1:80 | 1 |
| 2 |  | BUV395 | UV03 [420] (1.00) | CD40 | 5C3 | BD | 565202 | 1:40 | 2 |
| 3 |  | Autofluorescence | UV04 [440] (1.00) | NA | NA | NA | NA | NA | NA |
| 4 |  | BUV496 | UV07 [500] (1.00) | CD3 | UCHT1 | BD | 612940 | 1:40 | 2 |
| 5 |  | BUV563 | UV11 [575] (1.00) | CD56 | NCAM16.1 | BD | 565704 | 1:320 | 2 |
| 6 |  | Qdot 605 | UV13 [605] (1.00) | CD2 | S5.5 | Invitrogen | Q10172 | 1:160 | 1 |
| 7 |  | BUV615 | UV13 [605] (0.85),<br>YG03 [605] (1.00) | CD141 | 1A4 | BD | 624297 | 1:640 | 1 |
| 8 |  | Qdot 625 | UV14 [625] (1.00) | Streptavidin | NA | Invitrogen | A10196 | 1:200 | 2 |
| 9 |  | BUV661 | UV15 [655] (0.89),<br>R1 [655] (1.00) | CD303 | V24-785 | BD | 749920 | 1:80 | 1 |
| 10 |  | BUV737 | UV18 [725] (1.00) | CD86 | FUN-1 | BD | 612784 | 1:160 | 2 |
| 11 |  | BUV805 | UV21 [810] (0.65),<br>UV03 [420] (1.00) | CD45 | HI30 | BD | 564914 | 1:40 | 1 |
| 12 | 405 nm | BV421 | V01 [420] (1.00) | CD161 | DX12 | BD | 562615 | 1:20 | 1 |
| 13 |  | Super Bright 436 | V02 [440] (1.00) | IgM | SA-DA4 | Invitrogen | 62-9998- | 1:160 | 2 |
| 14 |  | Pacific Blue | V03 [460] (1.00) | CD1c* | L161 | Biolegend | 331508 | 1:80 | 2 |
| 15 |  | BV480 | V04 [475] (1.00) | CD28 | CD28.2 | BD | 566110 | 1:20 | 2 |
| 16 |  | AmCyan | V05 [500] (1.00) | CD19 | SJ25C1 | BD | 339190 | 1:20 | 1 |
| 17 |  | BV510 | V06 [515] (1.00) | CD11c | S-HCL-3 | Biolegend | 371513 | 1:320 | 1 |
| 18 |  | BV570 | V09 [575] (1.00) | CD45RO | UCHL1 | Biolegend | 304226 | 1:40 | 1 |
| 19 |  | BV605 | V11 [605] (1.00) | CCR7 | G043H7 | Biolegend | 353224 | 1:10 | 1 |
| 20 |  | BV650 | V13 [655] (1.00) | CD69 | FN50 | BD | 563835 | 1:40 | 1 |
| 21 |  | BV711 | V15 [700] (1.00) | FcER1 | AER-37 | Biolegend | 334638 | 1:80 | 1 |
| 22 |  | BV750 | V17 [750] (1.00) | CD103 | Ber-ACT8 | BD | 747099 | 1:320 | 2 |
| 23 |  | BV786 | V18 [780] (1.00) | CCR4 | 1G1 | BD | 744141 | 1:40 | 2 |
| 24 | 488nm | BB515 | B02 [515] (1.00) | CD25* | BC96 | BD | 567318 | 1:20 | 2 |
| 25 |  | RB545 | B04 [545] (1.00) | CD14 | M5E2 | BD | 569259 | 1:40 | 1 |
| 26 |  | NovaFluor Blue | B04 [545] (1.00) | CD8a | OKT8 | Invitrogen | H003T03 | 1:40 | 2 |
| 27 |  | NovaFluor Blue | B07 [605] (1.00) | CD4 | SK3 | Invitrogen | H001T03 | 1:20 | 2 |
| 28 |  | RB613 | B07 [605] (1.00) | BTLA (CD272) | J168-540 | BD | Custom | 1:20 | 2 |
| 29 |  | BB660-P2 | B09 [655] (0.53),<br>R01 [655] (1.00) | CD27 | M-T271 | BD | Custom | 1:640 | 2 |
| 30 |  | PerCP | B10 [675] (1.00) | CD11b | M1/70 | Biolegend | 101230 | 1:80 | 2 |
| 31 |  | BB700 | B11 [700] (0.90),<br>R03 [700] (1.00) | IgG | G18-145 | BD | 742235 | 1:80 | 2 |
| 32 |  | RB705 | B11 [700] (1.00) | TCRgd | B1 | BD | 13345- | 1:10 | 1 |
| 33 |  | RB744 | B13 [750] (1.00) | CD127 | HIL7R- | BD | 13345- | 1:80 | 2 |
| 34 |  | RB780 | B14 [780] (1.00) | TIGIT | TgMab-2 | BD | 569864 | 1:20 | 1 |
| 35 |  | PerCP-Fire 806 | B15 [810] (1.00) | IgD | W18340F | Biolegend | 307818 | 1:40 | 1 |
| 36 | 561 nm | PE | YG01 [575] (1.00) | CD57 | NK-1 | BD | 560844 | 1:500 | 2 |
| 37 |  | Spark YG 593 | YG03 [605] (1.00) | CD20 | 2H7 | Biolegend | 302368 | 1:80 | 2 |
| 38 |  | PE-Alexa Fluor | YG04 [625] (1.00) | CD24 | SN3 | Invitrogen | MHCD24 | 1:40 | 1 |
| 39 |  | PE/Fire 640 | YG05 [655] (1.00) | CXCR3 | G025H7 | Biolegend | 353764 | 1:20 | 2 |
| 40 |  | PE-Cy5 | YG06 [675] (1.00) | CD123 | 9F5 | BD | 551065 | 1:20 | 1 |
| 41 |  | PE-Cy5.5 | YG07 [700] (1.00) | ICOS (CD278) | ISA-3 | Invitrogen | 35-9948- | 1:40 | 2 |
| 42 |  | PE-Alexa Fluor | YG08 [725] (1.00) | CD16 | 3G8 | Invitrogen | MHCD16 | 1:320 | 2 |
| 43 |  | PE-Cy7 | YG10 [780] (1.00) | PD1 (CD279) | EH12.1 | BD | 561272 | 1:20 | 2 |
| 44 |  | PE/Fire 810 | YG11 [810] (1.00) | HLA-DR | L243 | Biolegend | 90000015 | 1:40 | 1 |
| 45 | 637 nm | APC | R01 [655] (1.00) | Va24-JA18 | 6B11 | Biolegend | 342908 | 1:40 | 1 |
| 46 |  | Alexa Fluor 647 | R02 [675] (1.00) | Va7.2 | 3C10 | Biolegend | 351726 | 1:80 | 1 |
| 47 |  | Spark NIR 685 | R03 [700] (1.00) | KLRG1 | SA231A2 | Biolegend | 367740 | 1:20 | 2 |
| 48 |  | R718 | R04 [725] (1.00) | CD39 | A1 | BD | 567674 | 1:160 | 2 |
| 49 |  | Zombie-NIR | R05 [750] (1.00) | Amine | NA | Biolegend | 423106 | 1:500 | NA |
| 50 |  | APC-Cy7 | R06 [780] (1.00) | CD163 | GHI/61 | Biolegend | 333622 | 1:80 | 1 |
| 51 |  | APC/Fire 810 | R07 [810] (1.00) | CD38 | HB-7 | Biolegend | 356643 | 1:640 | 2 |

1. Thaw cryopreserved PBMCs in 37°C water bath, slowly dilute in warm RP10, and wash twice in warm medium. Count cells. Use between 2-5 x 10^6^ cells per staining.
2. Resuspend cells in the appropriate volume and dispense into a 96-well plate.
3. Centrifuge plate at 400g for 5 minutes. Flick supernatant and dry the top of the plate by carefully placing it on a paper towel.
4. Add 100 ul of freshly prepared Fc-Block/Viability dye solution and incubate for 15 minutes at room temperature (RT) in the dark.
5. Wash by adding 200 µl of FACS buffer, centrifuge plate at 400g for 5 minutes, flick supernatant, and dry the top of the plate by carefully placing it on a paper towel.
6. During the last centrifugation of step 5, centrifuge antibody staining mix 1 at 10,000g for 5 minutes to pellet antibody aggregates. When pipetting in the following steps, avoid scraping the bottom of the tube so as not to disturb the pellet.
7. Add 50 ul of antibody staining mix 1 containing the correct final dilutions of all antibodies in Brilliant Stain Buffer Plus with True-Stain Monocyte Blocker. CellBlox Blocking Buffer is not needed during this stain as no NovaFluor reagents are within this staining mix. Incubate for 20 minutes at RT in the dark.
8. Add 200 ul of FACS buffer to all wells, centrifuge plate at 400g for 5 minutes, flick supernatant, and dry the top of the plate by carefully placing it on a paper towel. Repeat this wash once.
9. During the last centrifugation of step 8, centrifuge antibody staining mix 1 at 10,000g for 5 minutes to pellet antibody aggregates. When pipetting in the following steps, avoid scraping the bottom of the tube so as not to disturb the pellet.
10. Add 50 ul of antibody staining mix 2 or FMO staining mixes containing the correct final dilutions of all antibodies in Brilliant Stain Buffer Plus with True-Stain Monocyte Blocker and CellBlox Blocking Buffer. Incubate for 20 minutes at RT in the dark.
11. Add 200 ul of FACS buffer to all wells, centrifuge plate at 400g for 5 minutes, flick supernatant, and dry the top of the plate by carefully placing it on a paper towel. Repeat this wash once for a total of two washes.
12. Add 100 µl of Cytofix/Cytoperm to each well, resuspend, and incubate for 10 minutes at RT in the dark.
13. Add 200 µl of FACS buffer to all wells, centrifuge plate at 400g for 5 minutes, flick supernatant, and dry the top of the plate by carefully placing it on a paper towel.
14. Resuspend the cells in 100-200 ul of FACS buffer and keep wrapped in aluminum foil at 4°C until acquisition.

#### Preparing Single Stain Controls for Unmixing

1. In parallel to staining the full panel, prepare single stain controls for both beads and cells.
2. For single stain controls using cells, use 2-5 x 10^6^ cells per staining as in full the panel.
3. For only the Zombie-NIR viability control on cells, heat kill cells by incubating at 65 °C for 1 minute followed by placing on ice for 1 minute prior to staining. Heat killed cells should be divided in half. Stain one half with the Zombie-NIR viability dye and leave the other half unstained to use as the negative population for the Zombie-NIR parameter when unmixing.
4. Unstained cell controls will consist of: unstained cells, unstained cells treated with Brilliant Stain Buffer Plus, and unstained heat killed cells. Unstained cells are to be used for the negative population for all antibody based fluorescent probes. Unstained heat killed cell are to be used for the negative population for the viability dye Zombie-NIR. Unstained cells that were treated with Brilliant Stain Buffer are to be used to model autofluorescence in unmixing.
4. For single stain controls using beads, ¼ a droplet of the CompBeads Mouse (normal or plus) or Rat/Hamster was used per control respectively.
5. For the Zombie-NIR viability control on beads, divide one droplet of the positive ArC bead in half. Use half of the positive ArC bead to stain with the Zombie-NIR viability dye as a positive population and leave the other half unstained for the negative population when unmixing. We have observed differences in the autofluorescence of the positive and negative ArC beads, and therefore only use the positive ArC bead for the positive and negative populations of viability dyes.
6. Stain both cell and bead single stain controls at the same titer as described in **Online Table 2** in a 50 ul volume in FACS buffer.
7. Perform staining of single stain controls alongside staining of the full panel to ensure that controls are treated with the same buffers (PBS/FACS/ & Cytofix/Cytoperm) and for the same amount of time to best model experimental fluorescence.

